# Expanding High-Fidelity Multiplexing in Ultrasensitive Single-Molecule Protein Detection via Proximity Barcoding

**DOI:** 10.64898/2026.01.03.697469

**Authors:** Chi-Chia Wang, Emily Dorsey, Connie Wu

**Affiliations:** Life Sciences Institute, University of Michigan, 210 Washtenaw Avenue, Ann Arbor, MI 48109; Department of Biomedical Engineering, University of Michigan, Ann Arbor, MI 48109; Department of Pharmaceutical Sciences, University of Michigan, Ann Arbor, MI 48109

## Abstract

The human proteome presents a vast information reservoir for basic and diagnostics research, yet the low abundances of many proteins in biofluids pose an analytical challenge. While ultrasensitive methods such as digital enzyme-linked immunosorbent assay have expanded the window of detectable proteins, multiplexing with high accuracy, sensitivity, and throughput remains limited by cross-reactivity and signal readout channels To address this challenge, we introduce PRO-MOSAIX (PROximity-barcoded Molecular On-bead Signal Amplification for Individual MultipleXing), a high-accuracy, ultrasensitive multiplex digital immunoassay platform that integrates high-throughput single-molecule protein detection with DNA barcode-based proximity ligation. PRO-MOSAIX generates “ON” signals only from matched affinity reagents in proximity, minimizing false positives from cross-reactive binding. This approach overcomes the multiplexing ceiling imposed by fluorescence spectral overlap by employing a single signal readout channel and DNA barcoding. In conjunction, we further improve multiplexing accuracy by mitigating a secondary source of false positives from DNA-based signal amplification. As a proof of principle, we establish and validate a 15-plex PRO-MOSAIX assay in human plasma, with low femtomolar sensitivities and high measurement accuracies. PRO-MOSAIX is modular and utilizes common laboratory instrumentation with a high-throughput flow cytometric readout, providing a broadly accessible tool that bridges the gap between analytical sensitivity and high-order multiplexing.

## Introduction

The extraordinary complexity of the human proteome presents a wealth of biological information for understanding and diagnosing diseases. However, much of this potential remains untapped due to measurement technology limitations and fundamental gaps in protein biomarker identification. Human plasma alone contains thousands of proteins spanning over ten orders of magnitude in concentration^[1-2]^, but only a small fraction is routinely used in the clinic, in part due to challenges in identifying biomarkers with high sensitivities and specificities^[3-5]^. Many low abundance proteins in plasma and other biofluids remain inaccessible by traditional detection methods such as enzyme-linked immunosorbent assay (ELISA), which have low to sub-picomolar (pM; 10^-12^ M) detection limits. Another major challenge is the simultaneous detection of many proteins with both high sensitivity and accuracy. Such multiplexing capabilities are critical for capture of disease complexities and heterogeneities, measurement throughput and cost-effectiveness, and sample conservation.

Ultrasensitive detection methods such as digital ELISA have expanded the window of detectable proteins by three to four orders of magnitude over conventional ELISA in recent years, reaching low abundance biomarkers such as many cytokines, tumor-derived or tumor-enriched proteins, and neurodegenerative disease biomarkers^[6-10]^. In digital ELISA, single molecules are captured on a high excess number of antibody-coated paramagnetic beads, ensuring a Poisson distribution in which most beads carry one or zero molecules. Single beads are then isolated into individual femtoliter microwells or droplets for signal amplification from single molecules, enabling counting of “ON” and “OFF” beads. We recently developed a compartmentalization-free digital ELISA platform, Molecular On-bead Signal Amplification for Individual Counting (MOSAIC), that generates a fluorescent signal localized to each bead carrying a single target molecule via rolling circle amplification (RCA)^[9]^. This localized signal amplification enables high-throughput readout via flow cytometry and achieves low-to mid-attomolar (aM; 10^-18^ M) detection limits, enhancing sensitivity by an order of magnitude over conventional digital ELISA.

Despite advances in sensitivity, accurate multiplexing in digital ELISA remains severely restricted by cross-reactivity and limited signal output channels, with up to only eight-plex assays reported to date^[9, 11-12]^. Multiplexing is achieved using unique fluorescent dye-encoded capture beads for each analyte, but exponentially increasing combinatorial interactions between affinity reagents and analytes with increased multiplexing inevitably lead to cross-reactive binding and false-positive “ON” signals (**Figure 1A**). Cross-reactivity also poses a major barrier in other multiplex immunoassays^[13]^. Various strategies have been employed to mitigate cross-reactivity in multiplex immunoassays, including spatial or temporal separation of affinity reagents^[14-17]^ and proximity-based approaches requiring matched affinity reagent binding for signal generation^[18-20]^. However, achieving the ultrasensitive detection, multiplexing capacity, and throughput required to probe low-abundance proteins in biofluids with high accuracy and scalability, while maintaining cost-effectiveness and broad accessibility, remains difficult.

**Figure 1.**
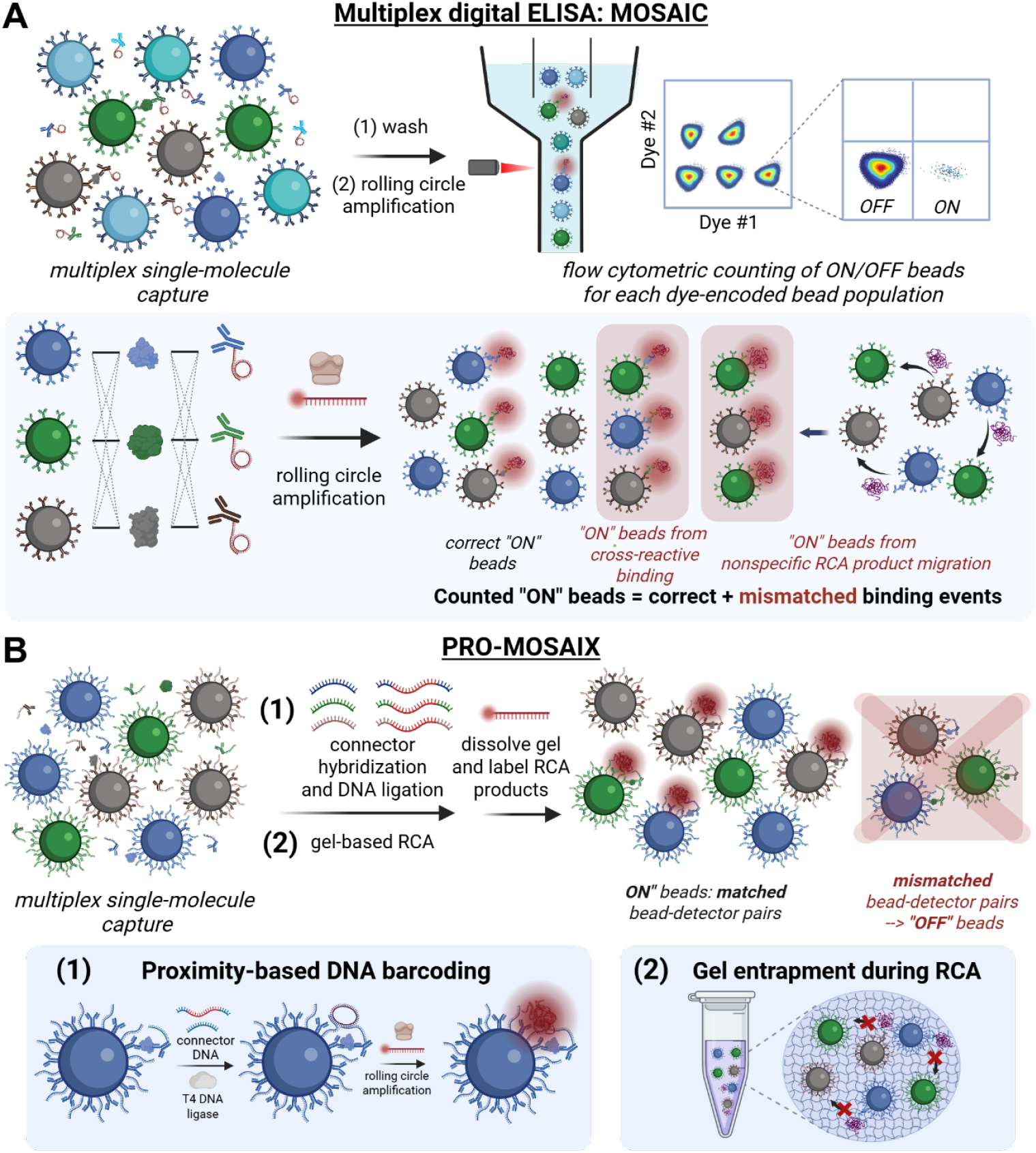
Multiplex digital immunoassay workflows and sources of cross-reactive signals. **(A)** Multiplex single-molecule protein detection with MOSAIC (Molecular On-bead Signal Amplification for Individual Counting). Single immunocomplex sandwiches are formed on a mixture of unique fluorescent dye-encoded beads, each coated with capture antibody for a specific analyte. Upon signal amplification via rolling circle amplification (RCA), fluorescent “ON” and “OFF” beads are counted for each bead population by flow cytometry. Cross-reactive interactions between different antibodies and analytes lead to mismatched binding events that undergo signal amplification, yielding incorrect “ON” beads. False positive “ON” beads can also result from nonspecific DNA product diffusion across beads during RCA. **(B)** Schematic of PRO-MOSAIX workflow. Each unique fluorescent dye-encoded bead is co-coupled with capture antibody and a proximity DNA barcode, while the detector antibody for each analyte is conjugated to a paired proximity DNA barcode. Upon multiplex single-molecule capture, only correctly formed single immunocomplex sandwiches will template the circularization of a DNA template after addition of a connector oligo mixture and DNA ligase (1: proximity-based barcoding). Each connector oligo set contains a shared sequence for hybridization of the same fluorescently labelled DNA probe after RCA. Nonspecific DNA product diffusion during signal amplification is prevented by encapsulating beads in an agarose gel during RCA (2: gel entrapment during RCA), after which the gel is dissolved for bead retrieval, labelling with fluorescent DNA probe, and flow cytometric counting.

Towards cross-reactivity mitigation in digital ELISA, a temporal separation strategy was previously developed, in which sequential sample incubation with each bead type and then corresponding detector antibody eliminated cross-reactive “ON” signals^[21]^. However, the increased assay time and workflow complexity with each additional analyte imposes a practical limit on the multiplexing order. More recently, we developed a barcoded MOSAIC platform, where each detector antibody is conjugated to a DNA template corresponding to a unique fluorescent DNA probe for RCA product labeling^[11]^. Correct capture-detector binding combinations are identified via matched bead and “ON” probe fluorescent colors, while mismatched color pairs from cross-reactive binding events are not counted. Although this strategy increased accuracy in multiplex digital ELISA, multiplexing capacity remains limited by the number of unique probe colors due to fluorescence spectral overlap. Breaking the multiplexing ceiling in digital ELISA thus remains a key challenge in improving the throughput and cost-effectiveness of low-abundance protein measurements.

Here, we introduce PRO-MOSAIX (**PRO**ximity-barcoded **M**olecular **O**n-bead **S**ignal **A**mplification for **I**ndividual Multiple**X**ing), a modular platform that enables high-order multiplexing in digital ELISA with minimal cross-reactivities. PRO-MOSAIX integrates the exceptional sensitivity, streamlined workflow, and broad accessibility of MOSAIC with the versatility of DNA barcoding and proximity ligation, where each capture bead and corresponding detector antibody are barcoded with a unique proximity oligo pair (**Figure 1B**). When in proximity in a correct immunocomplex sandwich, the matched DNA barcodes template the circularization and ligation of connector DNA oligos for RCA. Thus, only matched capture and detector reagents bound in a correct immunocomplex sandwich will generate an “ON” signal. Importantly, this approach uses a single signal output channel by encoding a common sequence for fluorescent DNA probe hybridization across all connector oligos, bypassing the multiplexing limit imposed by fluorescence spectral overlap. In conjunction, we identify an additional source of false positive signals in multiplex MOSAIC assays, arising from nonspecific DNA product diffusion between beads during signal amplification (**Figure 1A**). This phenomenon is addressed via reversible agarose gel encapsulation of beads during RCA to physically trap DNA products. Integrating the proximity barcoding and gel entrapment approaches vastly reduces cross-reactive signals in multiplex single-molecule protein detection. As a proof-of-principle, we developed and validated an ultrasensitive 15-plex PRO-MOSAIX assay with high measurement accuracies across low- and medium-abundance cytokine and tumor biomarkers in plasma. Our results establish PRO-MOSAIX as a high-fidelity, high-throughput multiplex digital immunoassay platform using common laboratory instrumentation, laying the foundation for massively multiplexed, ultrasensitive protein profiling across diagnostic and fundamental applications.

## Results and Discussion

### Development of Gel Entrapment Strategy to Mitigate Off-Target ON Signals During DNA Amplification

To inform our design of PRO-MOSAIX, we first investigated potential sources of false positive signals in addition to cross-reactive antibody-antigen binding in MOSAIC-based assays. As RCA and other DNA amplification methods can be prone to false positive signals arising from nonspecific amplification^[22]^, we explored this possibility by performing spike-in control experiments in which non-target fluorescent dye-encoded beads are added to beads with formed immunocomplex sandwiches labeled with circular DNA template, immediately prior to RCA (**Figure 2A**, top). We spiked in both antibody only-coupled beads used in MOSAIC and antibody/DNA-co-coupled beads that would be used in PRO-MOSAIX. Interestingly, we observed notable increases in the average molecules per bead (AMB) signals on the off-target beads after RCA, when spiked into a bead mixture carrying high numbers of DNA template-labeled immunocomplex sandwiches (**Figure S1**). This increase in false positive ON off-target beads occurred for both antibody-coupled and antibody/DNA-co-coupled beads. Such false positive signals may arise from nonspecific diffusion of DNA products across beads during RCA. While we were able to reduce these false positive signals by three-to five-fold via tuned RCA conditions with shorter reaction times and addition of low concentrations of heparin and salt, the off-target beads exhibited residual false positive signals from DNA product diffusion, with “cross-reactivities” of 0.5-0.6%, despite no exposure to analytes or detector antibodies (**Figure S1**).

**Figure 2.**
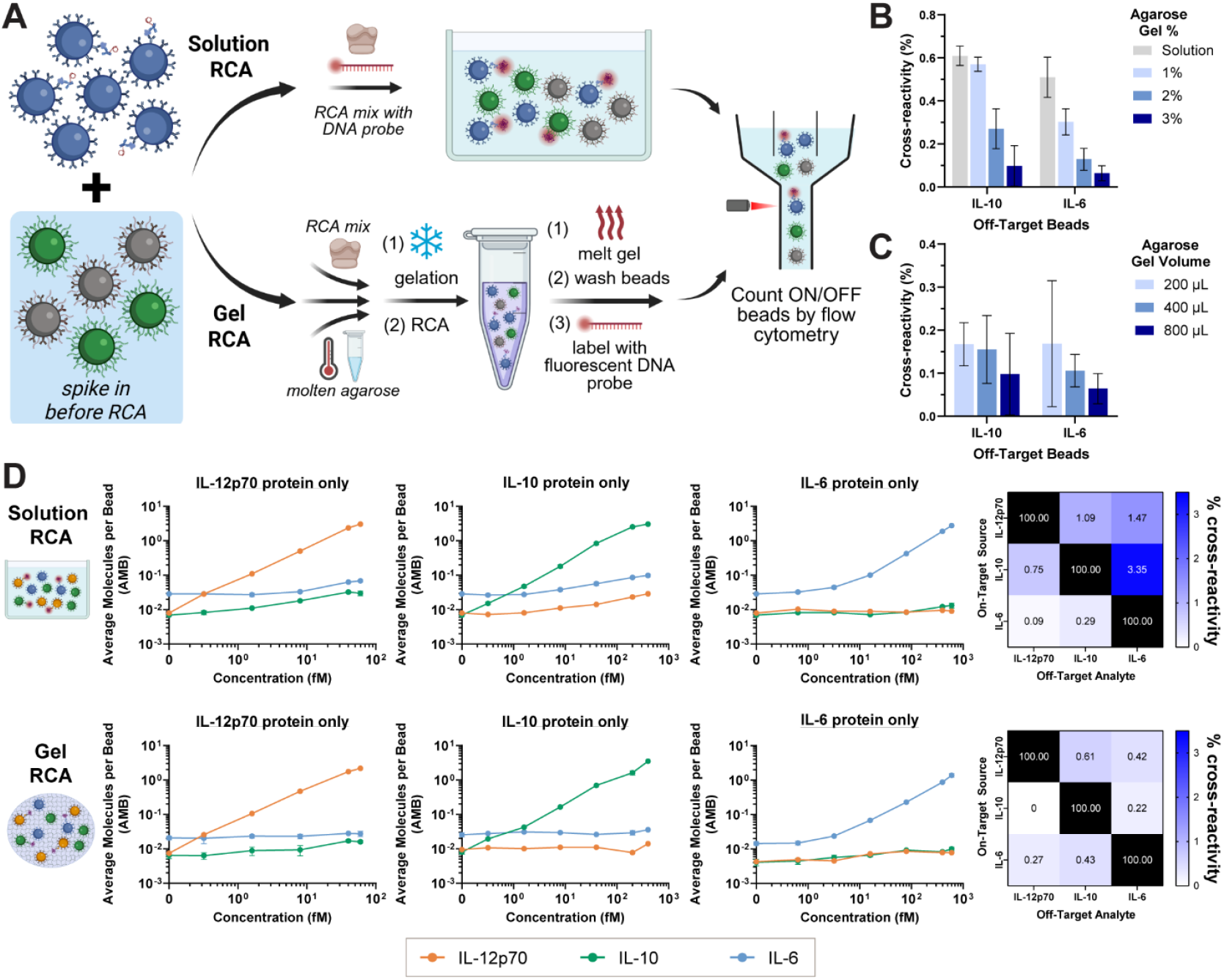
Identifying additional sources of false-positive signals arising from the DNA amplification process. **(A)** Schematic of spike-in controls in which off-target antibody/DNA co-coupled dye-encoded beads were added to beads carrying circular DNA template-labeled immunocomplexes, immediately before solution- or gel-based RCA. For solution-based RCA, beads were incubated with an RCA mixture containing phi29 polymerase and fluorescent DNA probe, and amplification proceeded in plates. For gel-based RCA, beads were transferred to tubes, resuspended in RCA mix without fluorescent probe, mixed with an equal volume of molten agarose, and solidified on ice; amplification occurred within the gel, after which beads were recovered by melting the gel at high temperature and labeled with probe. **(B-C)** Effects of agarose gel density **(B)** and volume **(C)** on false positive ON bead signals caused by nonspecific migration of DNA products during RCA. IL-12p70 beads (30,000 beads) and detector antibody were incubated with 30 fM IL-12p70, with 30,000 each of off-target IL-10 and IL-6 beads spiked in immediately prior to RCA for all conditions. Cross-reactivity (%) is defined as the ratio of off-target/on-target bead average molecules per bead (AMB) values after background subtraction. Error bars represent the standard deviation of three replicates. **(D)** Protein dropout curves across increasing concentrations of individual target proteins to evaluate cross-reactivities in a three-plex MOSAIC assay with solution- and gel-based RCA. Error bars represent the standard deviation of duplicate measurements, with the blank measured in triplicate. Cross-reactivity heatmap values were calculated from samples with on-target AMB values nearest 1.5 on each dropout curve and represent the mean of duplicate measurements.

We hypothesized that such nonspecific migration of DNA products can be inhibited by trapping the beads and DNA products in a reversibly formed hydrogel during RCA. In particular, we explored agarose gel encapsulation, as agarose gels are readily melted by heating for bead retrieval and have been successfully used with RCA in various applications.^[23-25]^ After non-target beads were spiked into beads carrying DNA template-labeled immunocomplexes, the beads were resuspended in RCA mixture, mixed with molten agarose, and immediately cooled on ice for gelation (**Figure 2A**, bottom). RCA was then carried out in the gel by incubating at 37°C, after which the gel was melted to retrieve beads for washing, fluorescent probe labeling of RCA products, and flow cytometric counting of ON and OFF beads. Compared to our optimized solution-based RCA, spatial separation of beads and entrapment of DNA products in an agarose gel successfully reduced false positive signals on off-target beads while maintaining high DNA amplification efficiencies (**Figure 2B** and **Figure S2A**). Increasing gel density up to 3% agarose decreased off-target bead ON signals, presumably via increased inhibition of DNA product diffusion.

Furthermore, as gel volume was increased from 200 µL to 800 µL for the same number of beads, cross-reactive signals from DNA product migration also slightly decreased together with improved consistencies, potentially due to greater bead separation (**Figure 2C**). Interestingly, on-target signal-to-background ratios also increased with increasing gel volume (**Figure S2B**). We therefore selected larger gel volumes for subsequent three-plex assays. To minimize potential nonspecific binding of any free DNA products during gel melting, we tested and optimized salt and heparin concentrations in the buffer that was added post-RCA to the gel prior to melting and used for subsequent bead washing (**Figure S3**). These results thus support our hypothesis that a dense hydrogel matrix can reduce or inhibit any nonspecific diffusion of large DNA products across spatially separated beads. Importantly, this method retains the streamlined solution-based readout of MOSAIC, does not require additional instrumentation or microfluidics, and is scalable for parallel processing.

Having verified the feasibility of our gel entrapment strategy for mitigating nonspecific DNA product diffusion across beads during signal amplification, we compared the cross-reactive signals of a full three-plex MOSAIC assay with solution-versus gel-based RCA. Human IL-12p70, IL-10, and IL-6 were used as model cytokine analytes that play important roles in inflammation, cancer, and infectious diseases^[26-28]^, with corresponding capture beads each encoded with a distinct AlexaFluor 488 dye intensity. Dropout curves, in which increasing concentrations of single protein analytes are measured with the multiplex assay, were performed to assess cross-reactivity. Notably, gel entrapment during RCA significantly decreased false positive signals on off-target beads at increasing on-target analyte concentrations (**Figure 2D**). Quantification of these false positive signals at high concentrations of each analyte yielded largely lower cross-reactivities across off-target beads using gel-based versus solution-based RCA (up to 0.61% and 3.35%, respectively). Our gel entrapment approach yielded sub-to low femtomolar limits of detection, which were slightly higher than the corresponding solution-based RCA format but remained comparable in range to other digital immunoassay platforms^[6, 10]^ (**Table 1** and **Figure S4**). This modest reduction in sensitivity may be attributed to lower bead recoveries from the gel, thereby decreasing the number of counted events and measurement precision at low concentrations.

**Table 1.**
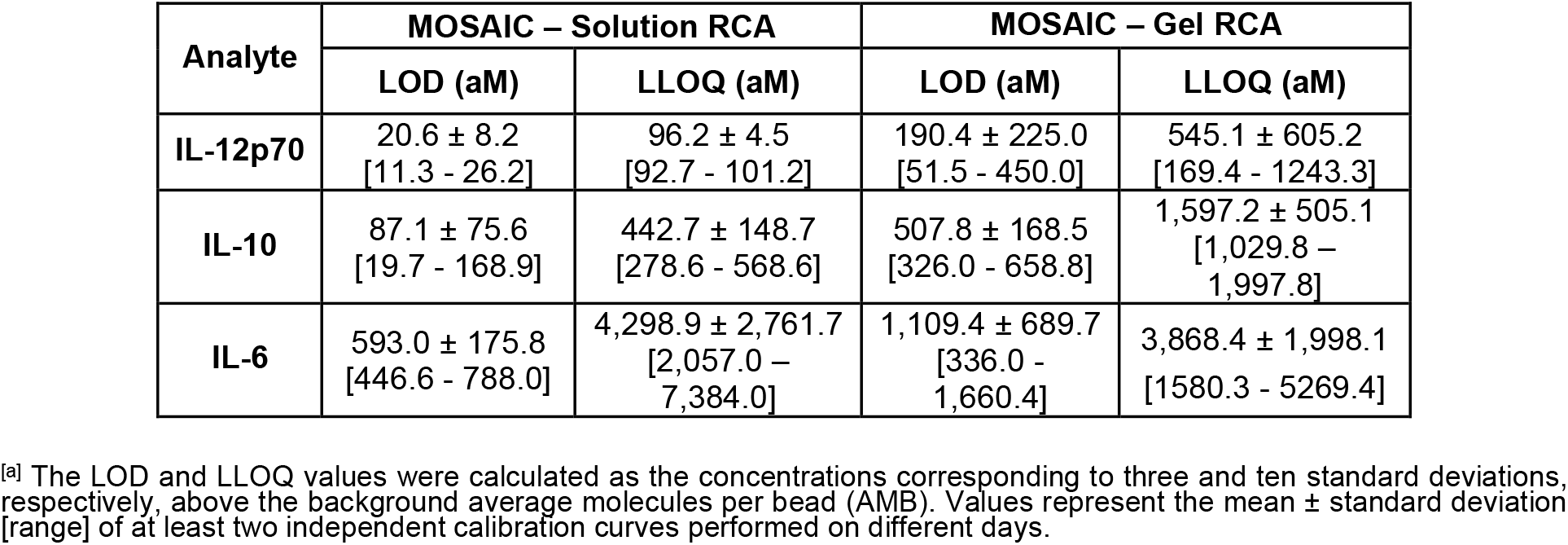
Limit of detection (LOD) and lower limit of quantification (LLOQ) values of three-plex MOSAIC assays with solution- and gel-based rolling circle amplification.^[a]^.

### Incorporation of Proximity-Based Barcoding for Accurate Multiplexed Single-Molecule Protein Detection

We next introduced proximity-based barcoding into the multiplex assay workflow, to develop the PRO-MOSAIX platform. For each of the analytes IL-12p70, IL-10, and IL-6, dye-encoded beads were co-coupled with capture antibody and a proximity oligo barcode, and detector antibody was conjugated with a paired proximity oligo barcode (**Figure 3A**). After single immunocomplex sandwiches are formed using a high excess number of beads over target molecules for each analyte, the capture and detector proximity DNA barcodes template the circularization and ligation of added connector DNA oligos. Connector oligos comprise hybridization sequences specific to each proximity barcode oligo pair and a common sequence for ATTO647N-DNA probe hybridization after RCA, thus requiring only one fluorescence channel to be used for “ON” bead signal readout. In parallel, the versatility of ratiometric fluorescent dye encoding of beads^[29-30]^ enables generation of a large array of capture beads that are distinguishable by flow cytometry.

**Figure 3.**
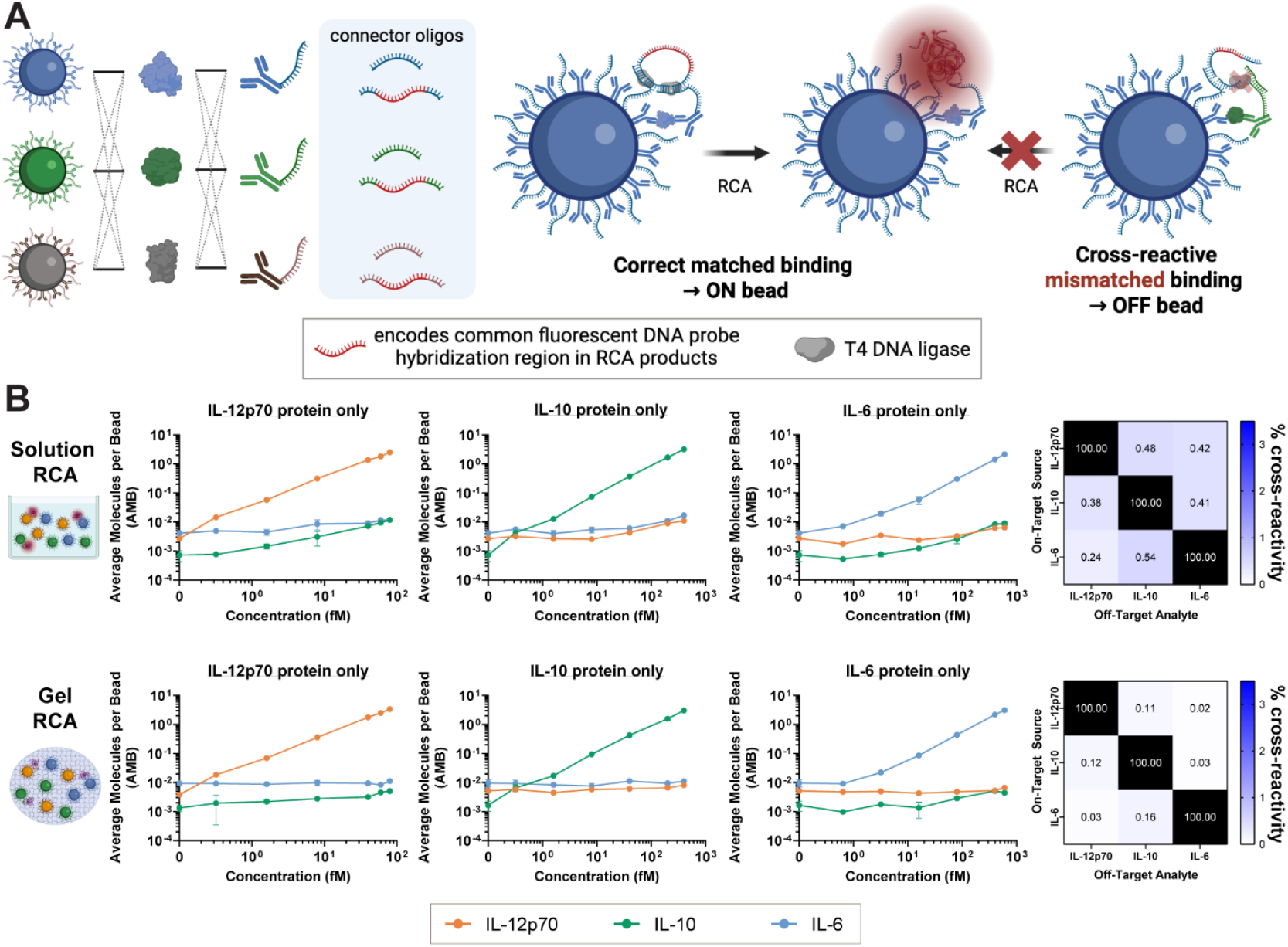
Proximity-based barcoding for multiplexed single-molecule protein detection. **(A)** Schematic of proximity ligation-based barcoding in PRO-MOSAIX. Dye-encoded beads are co-coupled with capture antibody and proximity DNA barcode, while detector antibodies are conjugated with a paired proximity DNA barcode. Once single immunocomplex sandwiches are formed, connector oligos comprising hybridization sequences specific for each proximity barcode pair and a common sequence (red) for fluorescent probe hybridization during RCA are hybridized and ligated into a circular template. Rolling circle amplification (RCA) is then carried out, generating a localized fluorescent “ON” signal on each bead carrying a correct immunocomplex sandwich. Signal is produced only when matched capture–detector pairs bind the same target molecule, enabling accurate high-order multiplexed detection through proximity barcoding. **(B)** Protein dropout curves across increasing concentrations of individual target proteins to evaluate cross-reactivities in a three-plex PRO-MOSAIX assay with solution- and gel-based RCA. Error bars represent the standard deviation of duplicate measurements, with the blank measured in triplicates. Cross-reactivity heatmap values were calculated from samples with on-target AMB values nearest 1.5 on each dropout curve and represent the mean of duplicate measurements. Cross-reactivity (%) is defined as the off-target/on-target average molecules per bead (AMB) ratio after background subtraction.

Integration of proximity barcoding maintained mid-aM to low fM sensitivities, with comparable detection limits as MOSAIC, for both solution- and gel-based RCA (**Table 2** and **Figure S5**). While proximity barcoding reduced relative cross-reactive signals in solution-based RCA format compared to the non-barcoded three-plex MOSAIC assay, dropout curves still exhibited increasing false positive signals at high single protein concentrations (**Figure 3B**). When we further incorporated gel entrapment during RCA, false positive signals were largely eliminated in the resultant three-plex PRO-MOSAIX assay, with minimal cross-reactivities (up to 0.16%). This trend was consistent with our previous observation of reduced cross-reactive signals in the three-plex MOSAIC assay with gel-based RCA, further supporting the efficacy of our gel entrapment strategy in mitigating false positive signals arising during DNA amplification. Notably, PRO-MOSAIX outperformed the gel-based non-barcoded MOSAIC assay in minimizing cross-reactivity, suggesting that proximity-dependent signal generation in our DNA-paired barcoding strategy can indeed reduce false positive ON bead signals from mismatched antibody-antigen binding.

**Table 2.** Limit of detection (LOD) and lower limit of quantification (LLOQ) values of three-plex PRO-MOSAIX assays with solution- and gel-based rolling circle amplification.^[a]^.

| Analyte | PRO-MOSAIX – Solution RCA |  | PRO-MOSAIX – Gel RCA |  |
| --- | --- | --- | --- | --- |
|  | LOD (aM) | LLOQ (aM) | LOD (aM) | LLOQ (aM) |
| **IL-12p70** | 25.2 ± 9.3<br>[18.7 - 31.8] | 109.1 ± 8.0<br>[103.4 - 114.8] | 31.1 ± 11.6<br>[17.1 - 43.1] | 119.3 ± 66.0<br>[49.6 - 182.5] |
| **IL-10** | 167.3 ± 101.9<br>[95.3 - 239.4] | 573.3 ± 240.1<br>[403.6 - 743.1] | 159.6 ± 112.2<br>[45.8 - 234.8] | 747.9 ± 598.8<br>[181.1 - 1,398.6] |
| **IL-6** | 518.7 ± 311.8<br>[298.2 - 739.1] | 1,741.1 ± 991.9<br>[1,039.7 - 2,442.4] | 941.8 ± 344.3<br>[488.1 - 1,273.6] | 3,604.6 ± 1,331.9<br>[1,660.3 - 4,681.0] |

<sup>[a]</sup> The LOD and LLOQ values were calculated as the concentrations corresponding to three and ten standard deviations, respectively, above the background average molecules per bead (AMB). Values represent the mean ± standard deviation [range] of at least two independent calibration curves performed on different days.

To further validate the accuracy of PRO-MOSAIX, we measured mixtures of varying high and low concentrations of the three protein analytes in the multiplex assay (**Figure 4**). Measurement accuracies were assessed by comparing the measured and actual concentrations of each protein and calculating percent “recovery.” For each assay, the low concentration of each analyte within a mixture was selected to be slightly above the lower limit of quantification, and the high concentration of each analyte was selected near the upper end of the corresponding calibration curve. Consistent with the high cross-reactivities observed in our dropout experiments, the three-plex MOSAIC assay with solution-based RCA yielded the least accurate measurements, with recoveries of approximately 200% for multiple low concentration analytes in the presence of high concentrations of other analytes. Incorporation of proximity barcoding while retaining the solution-based RCA format improved measurement accuracies for low concentration analytes but still resulted in overestimated measurements in one of the mixtures. When we further integrated our gel entrapment strategy, the resultant PRO-MOSAIX assay achieved high measurement accuracies, with 70-130% recoveries across all low concentration and nearly all high concentration analytes. For the three-plex MOSAIC assay, the gel entrapment strategy was also able to eliminate overestimation of low concentration analytes. These results suggest that much of the false positive signals in the three-plex assay arise during DNA signal amplification and are mitigated by our gel entrapment strategy during RCA. While cross-reactive antibody-antigen interactions are fewer in a three-plex assay compared to high-order multiplex assays, PRO-MOSAIX together with gel-based RCA yielded the most consistent reduction in cross-reactivities along with high measurement accuracies.

**Figure 4.**
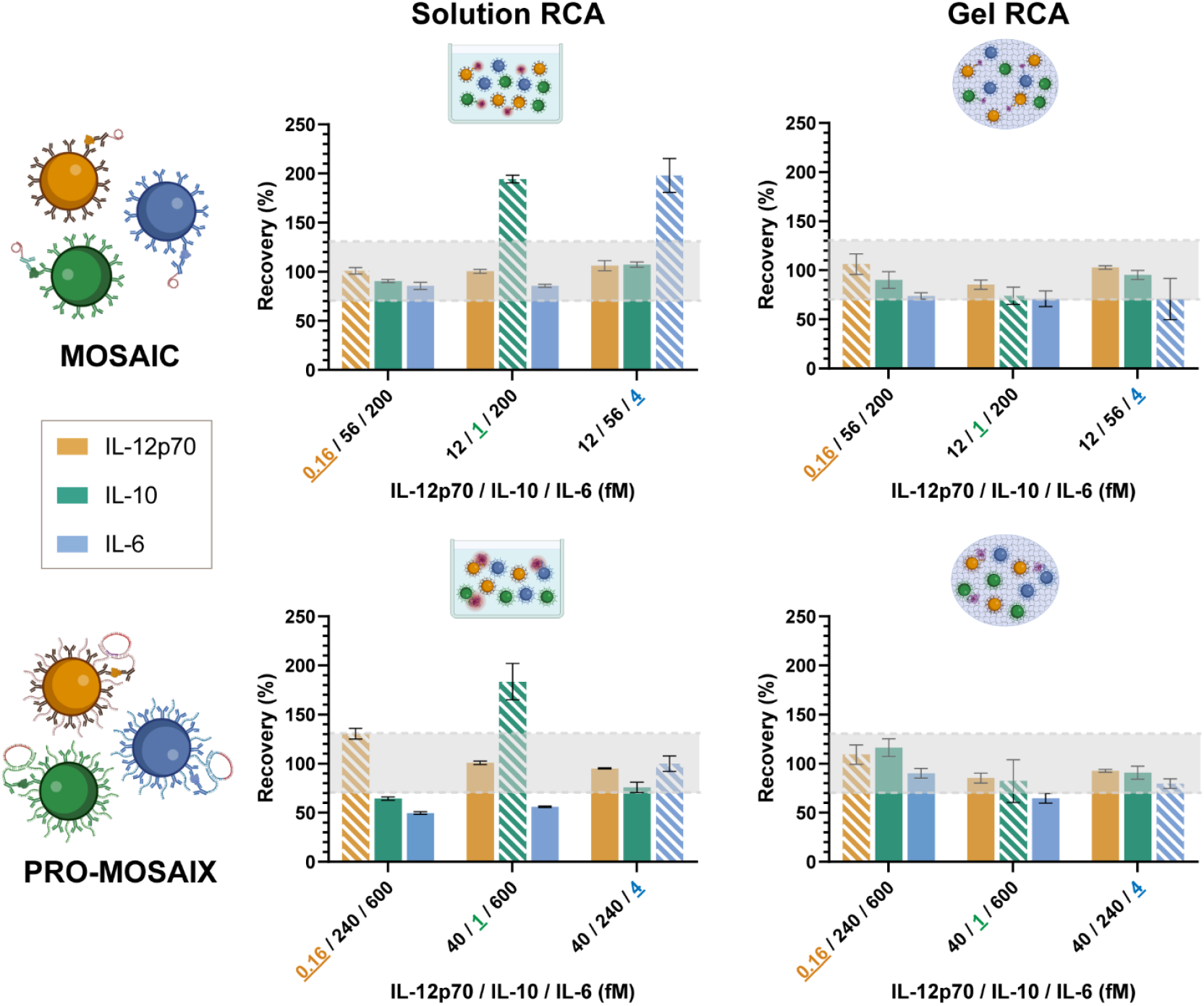
Measurement accuracies of three-plex MOSAIC and PRO-MOSAIX assays. Mixtures of recombinant proteins at varying high and low concentrations were measured with three-plex MOSAIC and PRO-MOSAIX assays, with solution- or gel-based RCA. Recoveries were calculated as measured concentration divided by actual concentration of each analyte, with the acceptable 70-130% range highlighted in gray. Low-concentration analytes within each mixture are denoted by striped bars, with low concentrations selected near the assay lower limit of quantification. High analyte concentrations were selected near the upper end of the assay dynamic range. Error bars represent the standard deviation of n=2-3 replicates.

### Development of an Ultrasensitive 15-plex PRO-MOSAIX Assay

To further assess the ability of proximity barcoding to minimize cross-reactivities from increasing mismatched antiboty-antigen interactions, we next expanded PRO-MOSAIX to a larger multiplex panel. While up to eight-plex digital ELISA assays have been demonstrated to date, increasing cross-reactive antibody-antigen interactions and fluorescence spectral overlap in signal readout have limited accuracy and multiplex order^[9, 11, 31]^. As PRO-MOSAIX allows the same fluorescence channel to be used for ON signals across all targets while providing a signal selection filter for matched affinity reagent pairs, we explored its high-order multiplexing capabilities in a proof-of-principle ultrasensitive 15-plex assay. To generate an array of 15 distinguishable capture beads, we labeled carboxylated paramagnetic beads with unique ratios of up to three fluorescent dyes (**Figure S6** and **Table S1**). Each encoded bead was subsequently co-coupled with a specific capture antibody and proximity DNA barcode. As target analytes, we chose both low- and medium-abundance blood biomarkers to test the versatility in dynamic range of PRO-MOSAIX. Our panel included an expanded set of cytokines that regulate immune and inflammatory responses in diverse diseases^[32-35]^, as well as the tumor biomarkers CA125 and HE4^[36-37]^.

With over three-fold higher total number of beads in the 15-plex assay (20,000 beads per target) compared to our previous three-plex assay (30,000 beads per target), we tested increased gel volumes for RCA. Bead spike-in experiments during RCA showed similar cumulative cross-reactivities across off-target beads from DNA product diffusion as gel volume was increased past our original 800 µL used for the three-plex assay (**Figure S7A**). However, we noted generally improved consistencies among replicates with higher gel volumes, as well as slightly increased signal-to-background for certain on-target beads when gel volume was increased up to 1,600 µL (**Figure S7B**). We therefore selected higher gel volumes for subsequent 15-plex experiments. In addition, due to the increased number of DNA oligos in the 15-plex PRO-MOSAIX assay, we used a thermotolerant DNA ligase, Hi-T4, at elevated ligation temperature to minimize potential false-positive signals from nonspecific hybridization and ligation of mismatched DNA barcodes. Performing the ligation step at an increased temperature of 45°C with Hi-T4 DNA ligase, together with splitting the DNA hybridization and ligation into separate steps, maintained similar on-target signal-to-background ratios (**Figure S8A**).

To evaluate cross-reactivity, we measured high-concentration single protein samples with the 15-plex PRO-MOSAIX and corresponding MOSAIC assays. Consistent with our three-plex assay results, PRO-MOSAIX integrated with gel-based RCA yielded minimal cross-reactivities (**Figure 5A**). In comparison, the corresponding 15-plex MOSAIC assay exhibited higher cross-reactivities of up to over 3%, suggesting that proximity-based barcoding effectively mitigates false positive ON signals from increasing combinatorial cross-reactive antibody-antigen interactions with higher-order multiplexing. Further comparison with our original hybridization and ligation conditions showed that elevation of ligation temperature with Hi-T4 DNA ligase indeed lowered overall cross-reactive signals in PRO-MOSAIX (**Figure S8B**). For both MOSAIC and PRO-MOSAIX, signal amplification in solution format yielded higher cross-reactivities compared to gel format, similar to our observations in the three-plex assays (**Figure S9**).

**Figure 5.**
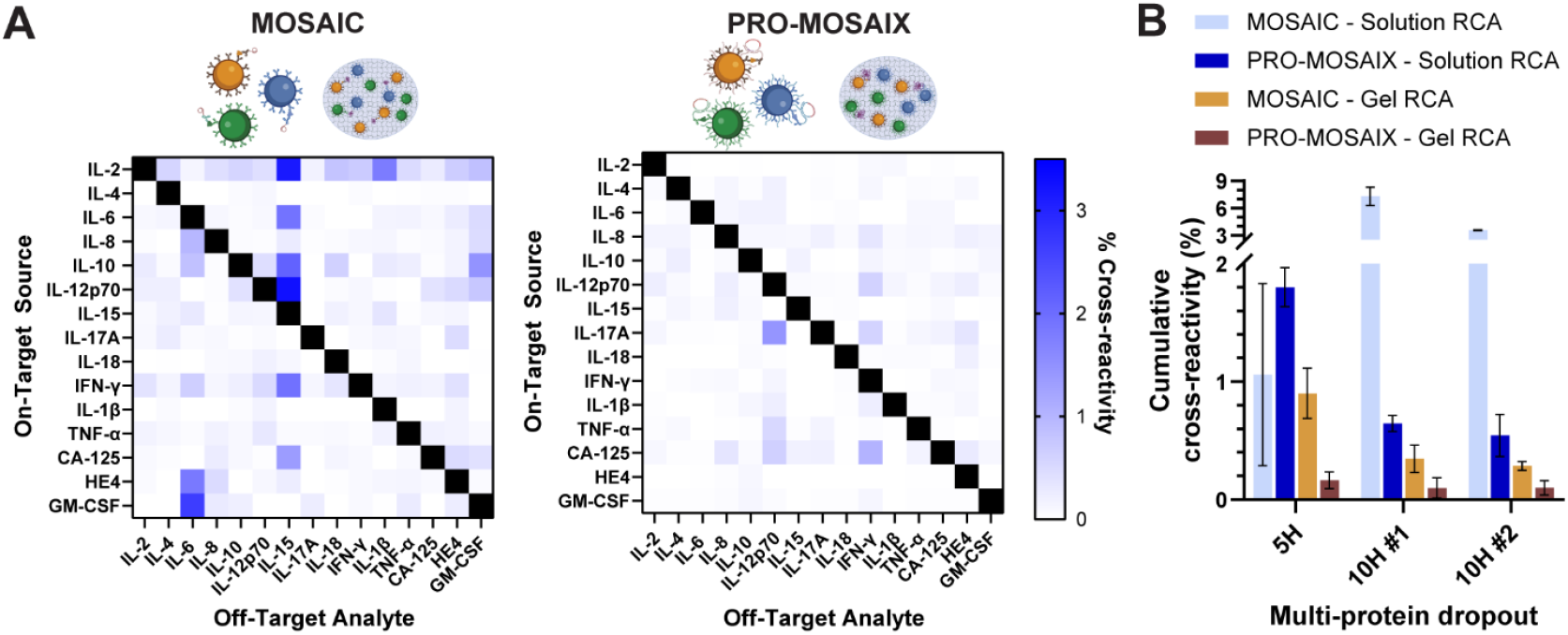
Cross-reactivity assessment in 15-plex MOSAIC and PRO-MOSAIX assays using single- and multi-protein dropout mixtures. **(A)** Heatmaps of percent cross-reactivity in 15-plex assays with gel-based RCA, in measurements of single high-concentration proteins. A gel volume of 2,400 µL used for RCA. Percent cross-reactivity is defined as the off-target/on-target average molecules per bead (AMB) ratio after background subtraction. The data represent the mean of duplicate measurements. **(B)** Cumulative cross-reactivities in 15-plex MOSAIC and PRO-MOSAIX assays in measurements of multi-protein dropout mixtures containing pooled high concentrations of five or ten proteins. A gel volume of 1,600 µL was used for RCA. Cumulative cross-reactivity is defined as the sum of off-target AMB values divided by the sum of on-target AMB values after background subtraction. Error bars represent the standard deviation of triplicate measurements. Mixture components are listed in **Table S2**.

When the 15-plex assays were further challenged with multi-protein dropout mixtures comprising pooled high concentrations of five or ten proteins, PRO-MOSAIX again exhibited much lower cross-reactivities on off-target beads compared to MOSAIC (**Figure 5B** and **Table S2**). With solution-based RCA, proximity barcoding substantially reduced but did not fully eliminate cumulative cross-reactivities across off-target beads in mixtures comprising ten high-concentration proteins. Similar to our single-protein dropout results, incorporating gel-based RCA generally decreased cumulative cross-reactivity for both MOSAIC and PRO-MOSAIC across different mixtures. Notably, combining proximity barcoding and gel-based RCA consistently yielded the lowest cumulative cross-reactivities across all mixtures. These trends were maintained across larger gel volumes, indicating robust suppression of cross-reactivity with our integrated approach (**Figure S10**).

We next assessed the analytical sensitivities of the 15-plex PRO-MOSAIX and MOSAIC assays. Notably, we observed significantly higher backgrounds and reduced dynamic ranges across multiple analytes in the 15-plex MOSAIC assay compared to PRO-MOSAIX (**Figures S11-S13**). Several analytes with particularly elevated backgrounds in MOSAIC, including IL-6, IL-15, and GM-CSF, exhibited less than two orders of magnitude in dynamic range and an order of magnitude lower sensitivity compared to PRO-MOSAIX (**Figure S13** and **Table S3**). Such a difference can be attributed to the cumulative nonspecific binding of all detector antibodies to each capture bead in multiplex MOSAIC. As every nonspecifically bound detector antibody can generate an ON signal, the background for each analyte increases with increasing multiplex panel size, which can reduce dynamic range and sensitivity. These elevated backgrounds and reduced dynamic ranges occurred in MOSAIC for solution-based RCA as well (**Figure S14** and **Table S4**). In contrast, the proximity barcoding strategy of PRO-MOSAIX restricts the background for each analyte in the multiplex assay to the nonspecific binding of only the corresponding detector antibody to the matched capture bead. With lower backgrounds and broader dynamic ranges, the 15-plex PRO-MOSAIX assay achieved sub-to low femtomolar limits of detection across most analytes.

### Validation of 15-plex PRO-MOSAIX Assay in Human Plasma

Finally, we evaluated the measurement accuracies of the 15-plex PRO-MOSAIX assay and its analytical performance in human plasma, using the gel RCA-integrated format. As the protein analytes span different concentration ranges in plasma, the assay dynamic range for each analyte can be readily tuned via detector antibody concentration and assay bead number^[9]^. We therefore decreased the detector antibody concentration for HE4, which can exist at mid-picomolar to low nanomolar concentrations in plasma, with particularly high levels in ovarian cancer patients^[38-39]^. The resultant 15-plex PRO-MOSAIX assay demonstrated sub-to high femtomolar detection limits, with a cumulative dynamic range of over six orders of magnitude across the 15 analytes (**Figure S15** and **Table S5**).

In mixed panels containing high- and low-concentration proteins, PRO-MOSAIX achieved high accuracies across nearly all low-concentration analytes in two distinct mixtures, yielding 70-130% recoveries even in the presence of ten high-concentration off-target proteins (**Figure 6A** and **Figure S16A**). When such mixtures were spiked into 12-fold diluted human pooled plasma, the 15-plex PRO-MOSAIX assay maintained high measurement accuracies for both low- and high-concentration spiked analytes (**Figure 6B** and **Figure S16B**). We further verified robust recoveries in mixtures containing all 15 recombinant proteins spiked at medium and high concentrations into diluted plasma (**Table S7**). Thus, the integrated PRO-MOSAIX platform achieves consistent measurement accuracies across low- and high-concentration analytes in various mixtures in both buffer and plasma.

**Figure 6.**
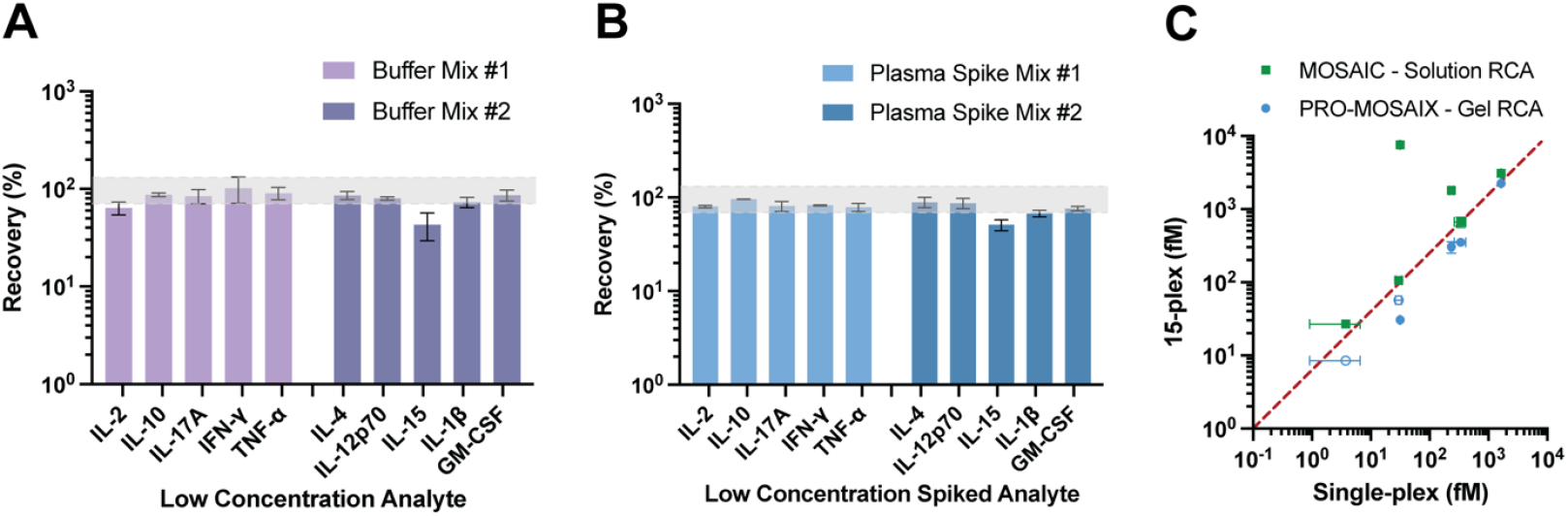
Measurement accuracies of 15-plex PRO-MOSAIX assay in protein mixtures and human plasma. **(A-B)** Recovery rates of 15-plex PRO-MOSAIX assay for low concentration analytes in recombinant protein mixtures in buffer **(A)** and spiked into 12-fold diluted pooled human plasma mixtures **(B)**. Each mixture in buffer or spiked into plasma comprised five low and ten high concentration protein analytes. Low concentrations were selected based on the assay lower limit of quantification or standard deviation of the measured endogenous concentration in plasma; high concentrations were selected based on the upper end of the assay dynamic range. A gel volume of 1,600 µL gel was used for RCA. Recoveries were calculated as the ratio of measured to actual analyte or spike concentrations, with the acceptable 70–130% range shaded in gray. Analyte concentrations are listed in **Table S6. (C)** Measured IL-4, IL-10, IL-12p70, IL-15, IL-17A, and CA125 concentrations in 12-fold diluted human pooled plasma using 15-plex MOSAIC (solution RCA) and PRO-MOSAIX (gel RCA) assays, compared to benchmark single-plex MOSAIC assays. Concentrations shown represent the endogenous plasma concentrations calculated from measured concentration values at 12-fold dilution. The red dashed line denotes perfect positive linear correlation between the multiplex and single-plex assays. The Pearson correlation coefficients were 0.1013 for the 15-plex MOSAIC (solution RCA) assay and 0.9987 for the PRO-MOSAIX (gel RCA) assay. Samples below the 15-plex assay LOD were assigned the LOD value and are marked as open squares or circles. Error bars represent the standard deviation of n=2-3 replicates.

To further validate the accuracy of the 15-plex PRO-MOSAIX in plasma, we compared the measured endogenous concentrations of several proteins in pooled human plasma with those obtained from corresponding single-plex MOSAIC assays (**Figure 6C**). PRO-MOSAIX measurements exhibited excellent agreement with the single-plex benchmarks across analytes spanning femtomolar to picomolar concentrations, with a Pearson correlation coefficient of 0.9987. In contrast, the 15-plex MOSAIC assay showed poor concordance with its single-plex counterparts, with a Pearson correlation coefficient of 0.1013. This discrepancy was largely driven by overestimated concentrations of certain analytes, including IL-4 and IL-17A, in the multiplex MOSAIC assay. Such overestimation is consistent with the expected accumulation of nonspecific detector antibody binding in larger multiplex MOSAIC panels, which is amplified in complex biofluids containing interfering matrix components that increase nonspecific binding. The paired proximity barcoding mechanism in PRO-MOSAIX bypasses this cumulative nonspecific binding problem, thereby minimizing inflated background signals and increasing measurement accuracy in biofluids. Collectively, these results demonstrate that the PRO-MOSAIX platform can simultaneously quantify endogenous proteins at femtomolar concentrations in human plasma with high accuracies comparable to single-plex MOSAIC assays.

## Conclusion

Proteins play critical roles in nearly all biological and pathological processes, but their vast informational and clinical potential remains unrealized due to technological gaps in protein detection. We have developed a modular, broadly accessible platform, PRO-MOSAIX, that breaks the multiplexing ceiling in digital ELISA, introducing expanded high-fidelity multiplexing capabilities to ultrasensitive single-molecule protein detection. By incorporating proximity ligation with paired DNA barcodes on matched affinity reagents, our platform eliminates false positive signals from cross-reactive binding interactions. Such a strategy enables a single fluorescence channel to be used for ON signals across all capture beads, greatly increasing multiplexing capacity while ensuring accuracy. We further enhance measurement accuracies by addressing nonspecific DNA product diffusion across beads during signal amplification, using a reversibly formed gel to physically trap such DNA products.

Our integrated platform achieves mid-attomolar to low femtomolar sensitivities across multiple analytes, demonstrating accurate measurements across various high- and low-concentration analyte mixtures and in plasma. The dynamic range for each analyte in the multiplex assay can be independently tuned via detector antibody concentration and capture bead number, allowing broad coverage across low and high abundance proteins in a single measurement. Furthermore, our proximity barcoding strategy markedly reduces background and yields broader assay dynamic ranges along with improved analytical sensitivities compared to conventional multiplex MOSAIC and other digital immunoassays, particularly as multiplex order increases. In complex biofluids, the cumulative nonspecific binding of detector affinity reagents in conventional multiplex digital immunoassays can be exacerbated by matrix components, further compromising measurement accuracies. Such potential interference is mitigated in PRO-MOSAIX, which confines background to the nonspecific binding of only the cognate detector affinity reagent. The solution-based flow cytometric readout further bypasses limitations on dynamic range and multiplex order faced by digital immunoassay platforms that rely on physical compartmentalization. Altogether, PRO-MOSAIX mitigates challenges of cross-reactivities and cumulative nonspecific binding that limit high-order multiplexing in ultrasensitive protein detection.

There remain several limitations in the current work and avenues for future investigation. While we have developed a high-accuracy 15-plex digital immunoassay, PRO-MOSAIX can be readily expanded to much higher multiplex order in future work due to the versatility of DNA barcoding. As ratiometric fluorescent dye labeling of beads, along with incorporation of varying bead sizes, can enable hundreds of distinguishable beads^[29-30, 40]^, we expect that optimization and expansion of the fluorescent dye labeling used in this work, along with exploration of additional dye conjugation methods, can yield similarly large arrays. Furthermore, while our 15-plex PRO-MOSAIX assay achieved sub-to low femtomolar detection limits, further improvements in sensitivity are required for robust detection of lower abundance analytes at attomolar or below concentrations. Towards this front, we anticipate that assay bead recoveries can be increased upon further optimization of gel dissolution and exploration of additional gel materials. As more beads are counted, increased sampling efficiencies reduce Poisson counting error and enable assay bead number tuning for maximizing signal-to-background ratios while maintaining measurement precision. In conjunction, screening and development of additional affinity reagents can be performed to enhance sensitivities as required for rare analytes. Importantly, proximity barcoding removes the requirement for extensive validation of affinity reagents in conventional multiplex assays for cross-reactivities, considerably increasing flexibility in affinity reagent screening and assembly of large multiplex panels.

Another limitation is the current throughput of our gel entrapment method. While the developed gel-based RCA method allows parallel tube processing during signal amplification and is integrated with an automated flow cytometric readout, the use of tubes for gel formation and dissolution reduces throughput compared to solution-based RCA in multi-well plates. Scalability can be increased in future work using deep-well plates or adjustable multichannel pipettes in appropriate setups. The sample processing steps of PRO-MOSAIX are also amenable to semi- or full automation with robotic liquid handlers. In conjunction, future work will explore additional reversible hydrogel materials to further tune gel mesh size and dissolution steps, to enable small gel volumes for high-throughput 96- or 384-well plate processing while ensuring robust inhibition of DNA product diffusion during signal amplification. Alternative hydrogel materials with bioreducible or pH-labile bonds can also bypass the need for heating steps in gel dissolution. However, for applications where most or all analytes in the multiplex assay have endogenous concentrations in the lower half of the calibration curve, proximity barcoding even without gel-based RCA can provide high measurement accuracies, as it minimizes false positive signals arising from cross-reactive binding interactions.

In summary, PRO-MOSAIX expands high-fidelity multiplexing capabilities in single-molecule protein detection while utilizing common laboratory infrastructure with a high-throughput signal readout. The proximity-dependent pairing of affinity reagents not only improves the analytical sensitivities and accuracies of multiplex digital immunoassays in biofluids, but also simplifies signal readout to a single fluorescence channel. PRO-MOSAIX thus provides a scalable path towards substantially expanding multiplex panel sizes without compromising assay performance. Its modular design, femtomolar and below sensitivities, and accessible workflow can potentially accelerate biomarker signature discovery, enable sample conservation, and maximize biological insight across basic and clinical applications.

## Supporting Information

The authors have cited additional references within the Supporting Information.

## Acknowledgements

This work was supported by the National Institutes of Health (R35GM154852 to C.W.), the Department of Defense Congressionally Directed Medical Research Programs (HT9425-24-1-0817 to C.W.), the University of Michigan Life Sciences Institute startup grant (C.W.), and the UM Biological Sciences Scholars Program (C.W.). The authors would like to thank Dr. Evan Alexander for many helpful discussions.

