## Supporting Information for "Expanding High-Fidelity Multiplexing in Ultrasensitive Single-Molecule Protein Detection via Proximity Barcoding"

### Materials and Methods

#### Materials

All DNA oligos were purchased from Integrated DNA Technologies, with sequences listed in **Tables S8-S9**. T4 DNA ligase, Hi-T4 DNA ligase, adenosine triphosphate (ATP), and deoxynucleotide triphosphates (dNTPs) were purchased from New England Biolabs, except phi29 DNA polymerase, which was purchased from Biosearch Technologies. All antibodies and recombinant proteins are listed in **Table S10**. Paramagnetic beads were purchased from Agilent (LodeStars 2.7 µm Carboxyl magnetic beads). Bioconjugation reagents were purchased from Thermo Fisher Scientific, MilliporeSigma, Vector Laboratories, and BroadPharm.

#### Preparation of Fluorescent Dye-Encoded Beads

Fluorescent dye-hydrazides were dissolved in either 1X PBS or dimethyl sulfoxide (DMSO) to a concentration of 10 mg/mL. $3.75\times10^8$ LodeStars paramagnetic beads were washed three times with 500 µL bead wash buffer (1x PBS with 0.1% Tween-20), two times with 500 µL conjugation buffer (50 mM MES, pH 6) and resuspended in the desired volume of conjugation buffer. Fluorescent dye hydrazide combinations used to make the capture beads for each analyte are shown in **Table S1**. A 1 mg vial of 1-ethyl-3-(3-dimethylaminopropyl carbodiimide hydrochloride (EDC, Thermo Fisher Scientific) was dissolved in 100 µL cold conjugation buffer. The desired volume of fluorescent dye hydrazide and 10 µL EDC were then added simultaneously to the beads. After shaking for one hour at room temperature, the resultant dyed beads were washed three times with 500 µL bead wash buffer, resuspended in 500 µL bead wash buffer, and shaken for an additional hour at room temperature. The beads were then resuspended in 500 µL 100mM sodium bicarbonate pH 9.3, shaken for one hour at room temperature, and resuspended in 400 µL storage buffer comprising 100mM sodium bicarbonate pH 9.3 with 0.02% sodium azide for storage at 4 °C. The dye-labeled beads were counted with a Beckman Coulter Multisizer 4e Coulter Counter prior to antibody coupling.

#### Preparation of Capture and Detector Reagents for MOSAIC Assays

***Capture antibody-coupled beads.*** Capture antibodies were first reconstituted or buffer exchanged with bead conjugation buffer (50 mM MES, pH 6.2). Buffer exchange was performed by centrifuging three times through a 30K Amicon Ultra-0.5 mL centrifugal filter (MilliporeSigma) at 14,000x*g* for five minutes, with the addition of bead conjugation buffer up to a total volume of 500 µL each time. The centrifugal filter was then inverted into a new collection tube and centrifuged at 1,000x*g* for two minutes to collect the buffer-exchanged antibody, followed by rinsing of the filter with additional buffer and another centrifuge cycle to recover more antibody.

The 2.7 µm carboxylated paramagnetic beads, either undyed or fluorescent dye-labeled were washed two times with 300 µL 10 mM NaOH solution, then three times with 300 µL bead wash buffer (1x PBS, 0.1% Tween-20), and two times with 300 µL bead conjugation buffer before resuspending in cold bead conjugation buffer. The carboxylic acid groups were then activated by adding the desired amount of 1-ethyl-3-(3-dimethylaminopropyl)carbodiimide) (EDC, no-weigh format, Thermo Fisher Scientific), freshly reconstituted to 10 mg/mL in cold bead conjugation buffer, and shaken at 4°C for 30 minutes (total volume of 200 µL). After EDC activation, the beads were washed once with 300 µL cold bead conjugation buffer before resuspension in the capture antibody solution. The beads were then shaken at 4°C for two hours for antibody conjugation, after which they were washed two times with 300 µL bead wash buffer and resuspended in PBS with 1% bovine serum albumin (BSA) and 1 mM EDTA for blocking. After shaking for 45 minutes at room temperature, the blocked beads were washed once with 300 µL bead wash buffer and once with bead storage buffer (50 mM Tris-HCl pH 7.5, 150 mM NaCl, 1% BSA, 0.1% Tween-20, 10 mM EDTA, 0.15% ProClin 300). Finally, the beads were resuspended in the bead storage buffer and stored at 4°C for further use. The antibody-coupled beads were counted with a Beckman Coulter Multisizer 4e Coulter Counter. Specific coupling conditions for each capture bead are listed in **Table S11**.

***DNA primer-circular template formation.*** The DNA primer-circular template to conjugate to detector antibody for signal amplification in MOSAIC was formed by first annealing a 5' azide-modified primer strand with the linear form of the template carrying a 5' phosphate group. A solution of 30 µM azide-modified primer and 30.3 µM template in NEBNext Quick Ligation reaction buffer (New England Biolabs) was heated at 95°C for two minutes and allowed to slowly cool to room temperature over 90 minutes. T4 DNA ligase was then added at a final concentration of 60 U/µL and the ligation reaction was incubated at room temperature for three hours. The ligated primer-template was then buffer exchanged with a 0.5 mL 7K MWCO Zeba spin desalting column (ThermoFisher Scientific) into 10 mM Tris-HCl (pH 7.5), 150 mM NaCl, and 1 mM EDTA and stored at -20°C.

***DNA template-conjugated detector antibodies.*** Detector antibody was reconstituted or buffer exchanged with 1x PBS. Buffer exchange was performed with a 30K or 50K Amicon centrifugal filter in the same process as described for capture antibody buffer exchange. The antibody was modified with dibenzocyclooctyne (DBCO) groups by adding 20-fold molar excess of DBCO-PEG4-N-hydroxysuccinimidyl ester (MilliporeSigma), which was freshly dissolved in DMSO to 200 mg/mL. The NHS conjugation reaction was incubated at room temperature for 30 minutes and then purified by five centrifugation cycles through a 30K Amicon centrifugal filter. The DBCO-modified antibody was then incubated with a two-fold molar excess of the azide-modified DNA primer-circular template overnight at 4°C. The final antibody-DNA template conjugate was diluted in a final storage buffer comprising 25 mM HEPES (pH 7.3), 150 mM NaCl, 0.5% BSA, 5 mM EDTA, and 0.02% sodium azide and stored in aliquots at -80°C.

**Preparation of Capture and Detector Reagents for PRO-MOSAIX Assays**

***Capture antibody/proximity DNA barcode co-coupled beads.*** Proximity DNA barcode with a 5' amine was first conjugated to methyltetrazine-PEG4-NHS ester (mTz-PEG4-NHS, Vector Laboratories) prior to bead coupling. DNA barcodes were first reconstituted with RNase-free water and then buffer exchanged with 50 mM sodium borate buffer, pH 8.5, using a 7K MWCO Zeba spin desalting column. The DNA barcode was then modified with methyltetrazine (mTz) groups by adding 25-fold molar excess of mTz-PEG4-NHS, which was freshly dissolved in DMSO to 20 mg/mL. The NHS conjugation reaction was incubated at room temperature for one hour and then purified by four consecutive 7K MWCO Zeba spin desalting columns into 10 mM Tris pH 7.5, 100 mM NaCl, and 1 mM EDTA.

For bead co-coupling, 2.7 µm carboxylated paramagnetic beads (undyed or fluorescent dye-labeled) were activated by EDC following the same protocol as for capture antibody-coupled beads used in MOSAIC assays. The EDC-activated beads were then co-conjugated with capture antibody and trans-cyclooctene-PEG6-amine (TCO-PEG6-amine, BroadPharm). TCO-PEG6-amine was first dissolved in DMSO to a concentration of 50 mg/mL and further diluted to 0.5 mg/mL with cold bead conjugation buffer. After the beads were washed with cold bead conjugation buffer following EDC activation, they were resuspended in 200 µL solution comprising capture antibody and 50 µL TCO-PEG6-amine (0.5 mg/mL), and the solution was shaken at 4°C for two hours for antibody/TCO co-conjugation. After two hours, beads were washed five times with 300 µL bead wash buffer and resuspended in bead coupling buffer (1x PBS with 800-1000 mM NaCl, 0.05% Tween 20, and 0.5 mM EDTA). Corresponding mTz-modified proximity barcode DNA and extra NaCl were added to the solution to a final concentration of 10 µM DNA barcode and 800-1000 mM NaCl. The beads were then shaken at room temperature for 1-1.5 hours for DNA conjugation. After DNA conjugation, the beads were washed two times with 300 µL bead wash buffer and blocked with 1% bovine serum albumin (BSA) and 1 mM EDTA in PBS following the same protocol as for capture antibody-coupled beads used in MOSAIC assays. Specific coupling conditions for each proximity-barcoded capture bead are listed in **Table S12**.

***Proximity DNA-conjugated detector antibodies.*** Detector antibody was conjugated to proximity DNA barcode following the same protocol as for DNA template-conjugated detector antibodies, using a 5' azide-modified DNA oligo.

**Multiplex MOSAIC Assays**

Multiplex MOSAIC assays were performed as previously described, using detector antibodies conjugated to DNA primer-circular template[1]. Briefly, mixtures of dye-encoded antibody-coated capture beads (10 µL) were incubated with 100 µL sample in a 96-well plate (Greiner Bio-One, 655096). For the three-plex assay, DNA template-conjugated detector antibody mixtures (10 µL) were incubated with the beads and sample simultaneously. Protein samples and conjugated detector antibodies were diluted in a sample diluent consisting of 1x PBS with 2% BSA, 5 mM EDTA, 0.1% Tween-20, 0.1 mg/mL heparin (MilliporeSigma, H3393), and 0.15% ProClin 300. After shaking at one hour at room temperature, the samples were washed with wash buffer (5x PBS, 0.1% Tween-20, 0.02% ProClin 300) using a Tecan HydroSpeed microplate washer. For the 15-plex assay, DNA template-conjugated detector antibody mixtures diluted in sample diluent (100 µL) were then incubated with the beads after initial target capture and washing to form single immunocomplex sandwiches, followed by another platewashing cycle. DNA oligos used in MOSAIC assays are listed in **Table S8** and assay conditions are listed in **Table S13**.

**Multiplex PRO-MOSAIX Assays**

Multiplex PRO-MOSAIX assays were performed following the same protocol as for multiplex MOSAIC assays in single immunocomplex sandwich formation, with proximity DNA/capture antibody-co-coupled beads and proximity DNA-conjugated detector antibodies. After single immunocomplex sandwich formation, connector oligos were added to a concentration of 5 nM in hybridization/ligation mix comprising 50 mM Tris pH 7.5, 10 mM $MgCl_2$, 0.25 mg/mL BSA, 1 mM ATP, and 0.8 U/µL DNA ligase. For the three-plex PRO-MOSAIX assay, connector oligo hybridization and ligation were combined in one step, which was performed at 37°C for 30 minutes with T4 DNA ligase. For the 15-plex PRO-MOSAIX assay, the DNA hybridization and ligation steps were carried out sequentially, with hybridization performed at 37°C and ligation at 45°C with Hi-T4 DNA ligase, respectively, each for 30 minutes with shaking. Specific connector oligo pairs for each analyte are listed in **Table S9** and PRO-MOSAIX assay conditions are listed in **Table S14**. After connector DNA hybridization and ligation, the beads were washed with another platewashing cycle prior to bead transfer for subsequent signal amplification. Solution- and gel-based RCA were carried out as described below.

***Solution-based signal amplification.*** For solution-based RCA in multiplex MOSAIC assays, beads were transferred to a new plate (Greiner Bio-One, 655901), washed once with 1x PBS with 0.1% Tween-20, and resuspended in 50 µL RCA reaction mixture comprising 0.1 U/µL phi29 DNA polymerase (Biosearch Technologies), 0.25 mM deoxynucleotide solution mix, 2 mg/mL BSA, 0.2% Tween-20, 0.1% trehalose, 0.005 µg/mL heparin, and 2 nM ATTO 647N-labeled DNA probe in 1X reaction buffer comprising 50 mM Tris-HCl (pH 7.5), 10 mM $MgCl_2$, 10 mM $(NH_4)_2SO_4$, and 50 mM NaCl. The plate was shaken at 37°C for 20 minutes, after which the RCA reaction was quenched by adding 160 µL stop buffer comprising 1x PBS with 10 mM EDTA, 0.1% Tween-20, and 0.05% BSA. The samples were washed twice with 150 µL stop buffer and resuspended in the same stop buffer for flow cytometry. Samples were analyzed on a NovoCyte Advanteon flow cytometer (Agilent) equipped with three lasers. The specific RCA conditions of each assay are detailed in **Table S15**.

***Gel-based signal amplification.***
***Bead Preparation.*** Following the final plate wash after immunocomplex sandwich formation, beads were transferred from each well into 2-mL Protein LoBind tubes (Eppendorf) by gentle pipetting. The beads were washed once with PBS containing 0.1% Tween-20 to remove residual reagents prior to rolling circle amplification (RCA).

***Agarose gel-based RCA.*** RCA reactions were prepared in solutions containing 0.1 U/µL phi29 DNA polymerase, 0.2 mg/mL BSA, 0.2% Tween-20, and 0.1% trehalose in 1X reaction buffer (50 mM Tris-HCl (pH 7.5), 10 mM $MgCl_2$, 10 mM $(NH_4)_2SO_4$). For 3-plex assays, 400 µL of ice-cold RCA mix was added to bead pellets; for 15-plex assays, 800 or 1,200 µL was used. Beads were thoroughly resuspended and shaken on a Thermomixer C (Eppendorf) at 4°C.

Separately, equal volumes of 6% low-melting agarose aliquots (Fisher Scientific, BP165) were melted at 95 °C for two minutes, supplemented with dNTPs to a final concentration of 0.25 mM, and cooled at 50 °C for four minutes. The RCA-bead mixture was then combined with warm agarose in a single addition and vortexed immediately to form a uniform 3% agarose mixture. Final RCA reaction gel volumes were 800 µL for 3-plex assays and 1,600 or 2,400 µL for 15-plex assays. Agarose-RCA mixtures were gelled on ice for two minutes and subsequently incubated at 37 °C for 90 minutes for RCA.

Following RCA, gels were processed to recover beads from agarose. Tubes were placed on ice and supplemented with gel melting stop buffer (400 µL for 3-plex assays; 800 or 1,600 µL for 15-plex assays, comprising 1x PBS with 45 mM EDTA, 0.2% Tween-20, 0.2% BSA, and 0.15 mg/mL heparin). Samples were heated at 95 °C for 2-3 minutes on a ThermoMixer C (3-plex assays) or 5-6 minutes in a preheated water-filled rack (15-plex assays) to fully melt the agarose. Liquefied agarose was removed after two minutes of magnetic bead separation.

The recovered beads were then washed four times with stop buffer comprising 1x PBS with 10 mM EDTA, 0.1% Tween-20, 0.05% BSA, and 0.05 mg/ml heparin. During the first wash (600 µL), the beads were reheated at 95 °C for one minute to ensure complete melting and removal of any residual agarose, and then cooled to room temperature before additional washes. Subsequent washes used 600 µL, 400 µL, and 180 µL of prewarmed (80°C) stop buffer, with 2-minute magnetic separation between washes. For 15-plex assays, larger volumes of stop buffer were used for the four washes (1000 µL, 1000 µL, 1000 µL, and 180 µL), with the first wash containing 10% Agarose Dissolving Buffer (Zymo Research). After the final wash, beads were transferred to a 96-well plate (Greiner Bio-One, 655096) for DNA probe hybridization, using an adjustable multichannel pipette.

***Probe Hybridization and Detection.*** For post-amplification labeling, beads were washed once with probe hybridization buffer without probe (10 mM Tris pH 7.5, 10 mM $MgCl_2$, 0.05% Tween-20, and 0.005 μg/ml heparin), then incubated in 50 μL of the same hybridization buffer containing 10 nM ATTO 647N-labeled DNA probe. Hybridization was performed at 37 °C for 20 minutes with shaking. The beads were then washed once and resuspended in stop buffer for analysis on a Novocyte Advanteon flow cytometer.

**Plasma Samples**

Human plasma (K2 EDTA) samples (Bioreclamation IVT) were centrifuged at 2,000x*g* for 10 minutes at 4°C to remove cellular debris before diluting 12-fold in sample diluent for measurements. All plasma samples were de-identified and experiments were performed under Institutional Review Board approval by the University of Michigan.

**Data Analysis**

***Flow cytometric analysis.*** Flow cytometry data were analyzed with FlowJo Software (BD Biosciences) or NovoExpress Software (Agilent Technologies). After gating single beads by forward and side scatter, dye-encoded bead populations were distinguished via the appropriate fluorescence channels, with the same gates used across corresponding MOSAIC and PRO-MOSAIX assays with the same bead dye-labeling schemes. Representative gates for the 15-plex assay are shown in **Figure S6**. For each bead population in each assay, "ON" and "OFF" beads were determined by setting a fluorescence intensity threshold where less than 0.02-0.1% of the beads were "ON" in control samples comprising beads incubated with RCA mixture without phi29 (for solution RCA) or ATTO647N-DNA probe labeling solution (for gel RCA). The threshold was determined for each bead type in each experiment and applied to all samples. The fraction of "ON" beads was then converted to an AMB (Average Molecules per Bead) value according to the Poisson distribution.

***Calibration curves.*** Calibration curves for all analytes were generated by fitting AMB values to a four-parameter logistic (4PL) regression model using GraphPad Prism. The limit of detection (LOD) and lower limit of quantification (LLOQ) values were calculated as the concentrations corresponding to three and ten standard deviations, respectively, above the background AMB.

***Cross-reactivity analysis.*** For single protein samples measured with multiplex assays, percent cross-reactivities were calculated as the ratio of the off- to on-target bead (signal AMB - background AMB) values, where signal corresponds to the high-concentration single protein sample and background corresponds to a buffer-only sample. Cumulative cross-reactivities in the multi-protein dropout samples measured with the 15-plex assays were calculated as the ratio of the sum of all off-target bead (signal AMB - background AMB) values to the sum of all on-target bead (signal AMB - background AMB) values.

***Outlier identification.*** Outliers were identified using the two-sided Grubbs' test in GraphPad Prism with alpha value = 0.05. Critical Z-value is set to be no larger than 1.155 for triplicate measurements.

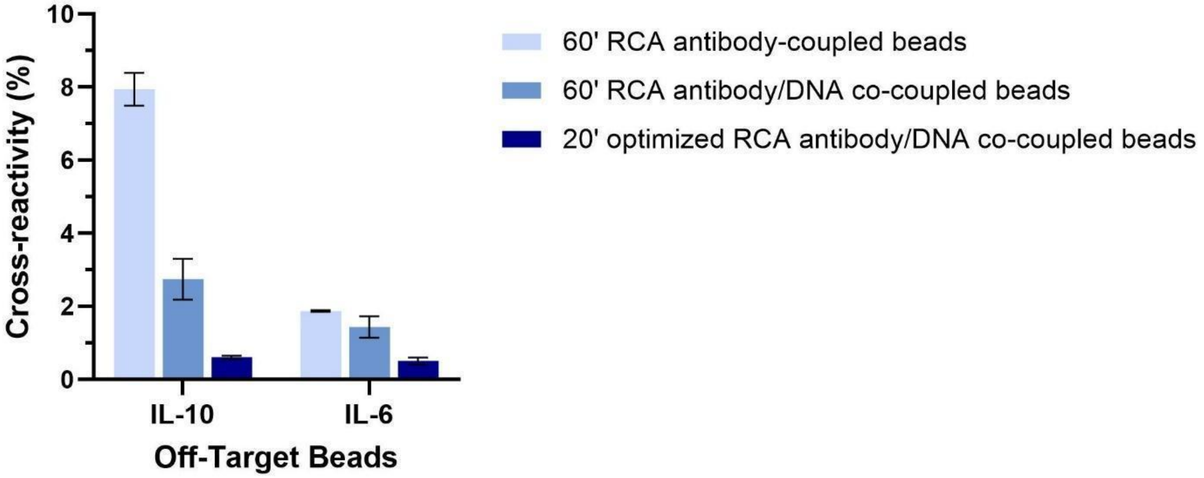


**Figure S1. False positive ON signal generation from nonspecific DNA product migration during solution RCA in MOSAIC.** False positive signals during RCA, calculated as percent cross-reactivities, on off-target antibody- or antibody/DNA-coupled beads spiked into a mixture of IL-12p70 capture beads carrying high numbers of DNA template-labeled immunocomplex sandwiches immediately before rolling circle amplification (RCA). Percent cross-reactivities were calculated via the ratio of off- to on-target bead [signal - background AMB] values. Optimization of RCA conditions with shorter amplification time and addition of 0.005 μg/mL heparin and 50 mM NaCl decreased cross-reactive signals arising from nonspecific DNA products during RCA. Error bars represent the standard deviation of triplicate measurements. The RCA conditions are listed in **Table S15**.

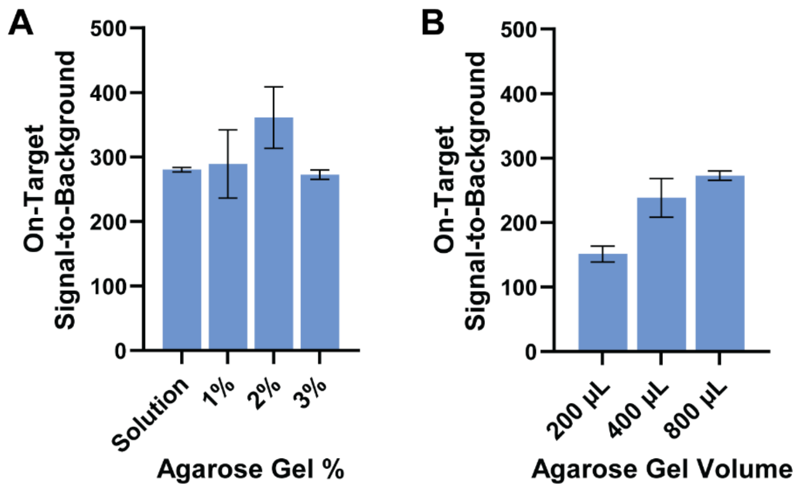


**Figure S2. On-target bead signal-to-background ratios with agarose gel entrapment during RCA.** Signal-to-background ratios for on-target IL-12p70 beads in three-plex spike-in RCA experiments, for **(A)** varying agarose gel percentages; and **(B)** varying agarose gel volumes. IL-12p70 beads (30,000 beads) and detector antibody were incubated with 30 fM IL-12p70 for all gel parameters, with 30,000 each of off-target IL-10 and IL-6 beads spiked in immediately before RCA. Solution RCA in **(A)** represents a 60-minute RCA reaction performed in solution in a 96-well plate. Error bars represent the standard deviation of triplicate measurements.

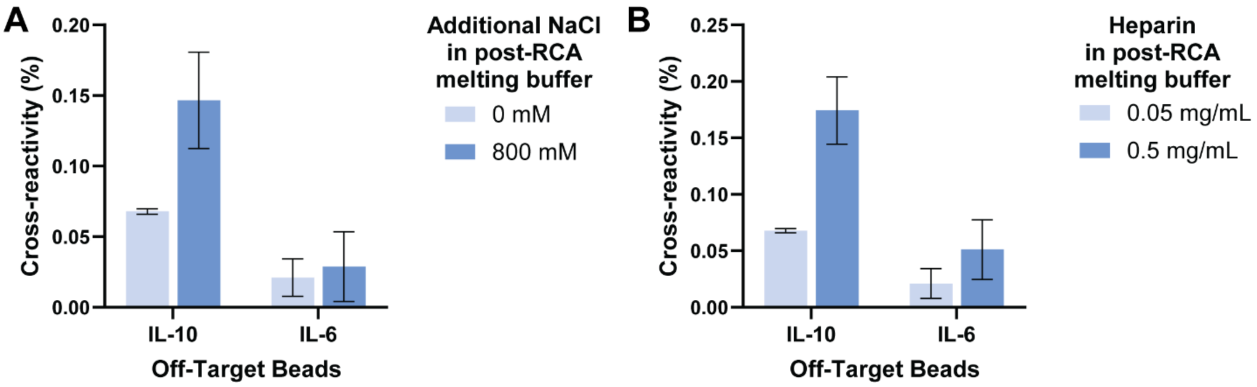


**Figure S3. Additional experimental parameters optimized in gel entrapment method.** Effects of additional salt **(A)** and heparin **(B)** in the post-RCA gel melting stop buffer on false positive signals, calculated as percent cross-reactivities, on off-target antibody/DNA-coupled beads spiked into a mixture of IL-12p70 capture beads carrying high numbers of DNA template-labeled immunocomplex sandwiches immediately before gel-based RCA. Percent cross-reactivities were calculated via the ratio of off- to on-target bead (signal AMB - background AMB) values. Error bars represent the standard deviation of duplicate measurements.

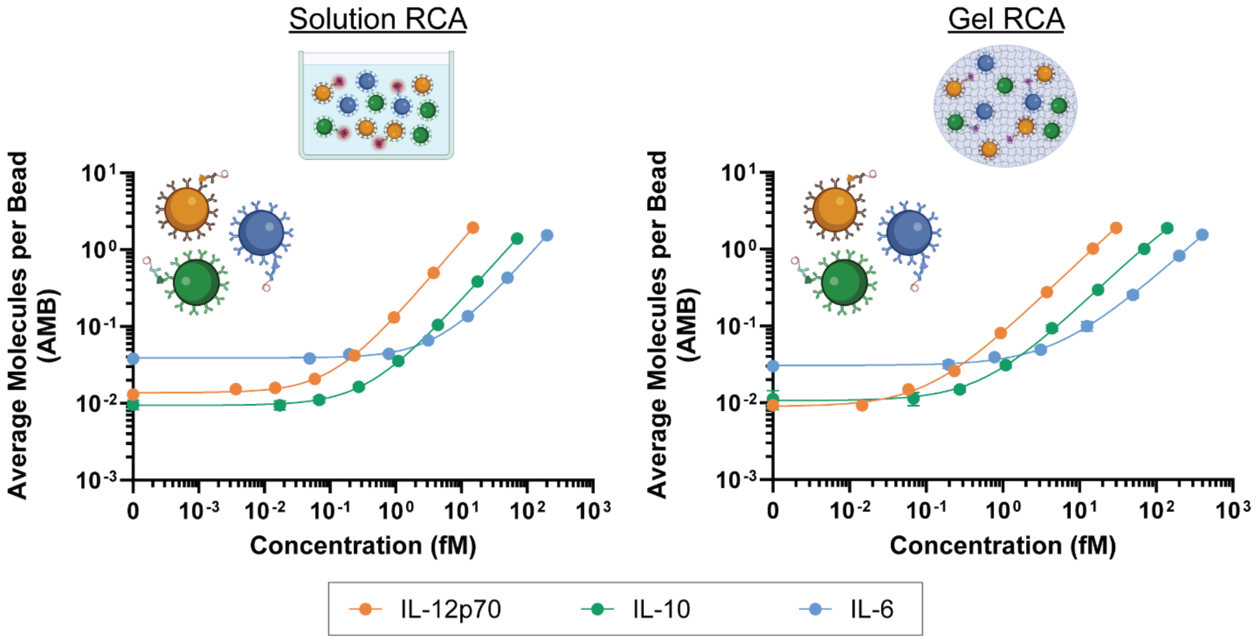


**Figure S4. Representative calibration curves for three-plex MOSAIC assays performed with solution- and gel-based RCA.** Calibration curves were fitted with four-parameter logistic (4PL) regression. Error bars represent the standard deviation of triplicate measurements, with n=4 replicates for the blank. The average LODs and LLOQs are summarized in **Table 1**.

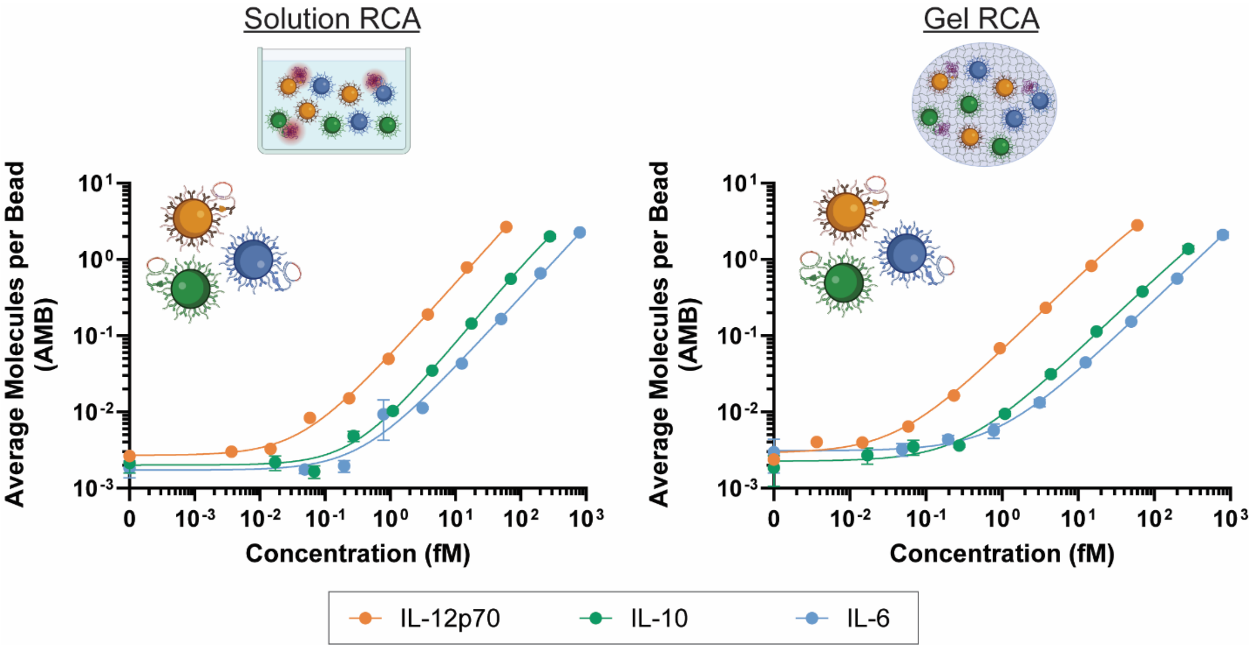


**Figure S5. Representative calibration curves for three-plex PRO-MOSAIX assays performed with solution- and gel-based RCA.** Calibration curves were fitted with four-parameter logistic (4PL) regression. Error bars represent the standard deviation of triplicate measurements, with n=4 replicates for the blank. The average LODs and LLOQs are summarized in **Table 2**.

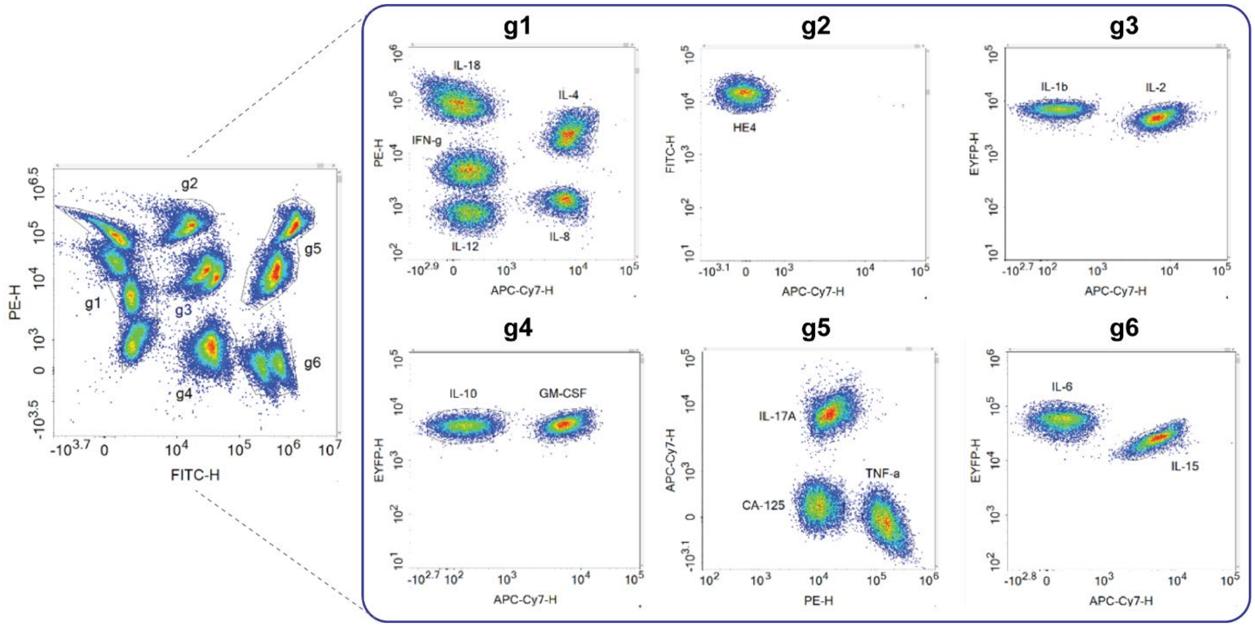


**Figure S6. Flow cytometry gating strategy for 15-plex PRO-MOSAIX assay.** Representative screenshots of 15 gated bead populations. Each bead type was labeled with a combination of CF 488A, BDP 558/568, and/or Cy7 hydrazides, as detailed in **Table S1**.

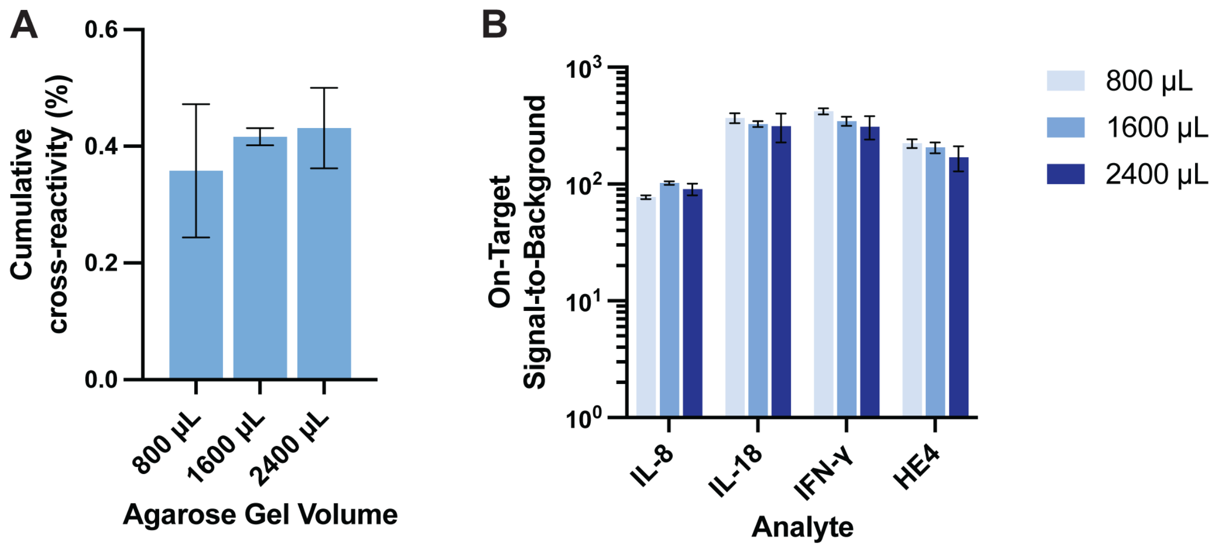


**Figure S7. Gel-volume optimization for the 15-plex PRO-MOSAIX assay. (A)** Cumulative cross-reactivity and **(B)** signal-to-background ratios of on-target beads in an antibody/DNA–coupled beads spike-in experiment. Off-target beads were added to a four-protein mixture before RCA using 800, 1600, or 2400 µL gel volumes. Percentage of cumulative cross-reactivity is defined as the sum of off-target signals divided by the sum of on-target signals after background subtraction. Error bars represent standard deviations of three replicates.

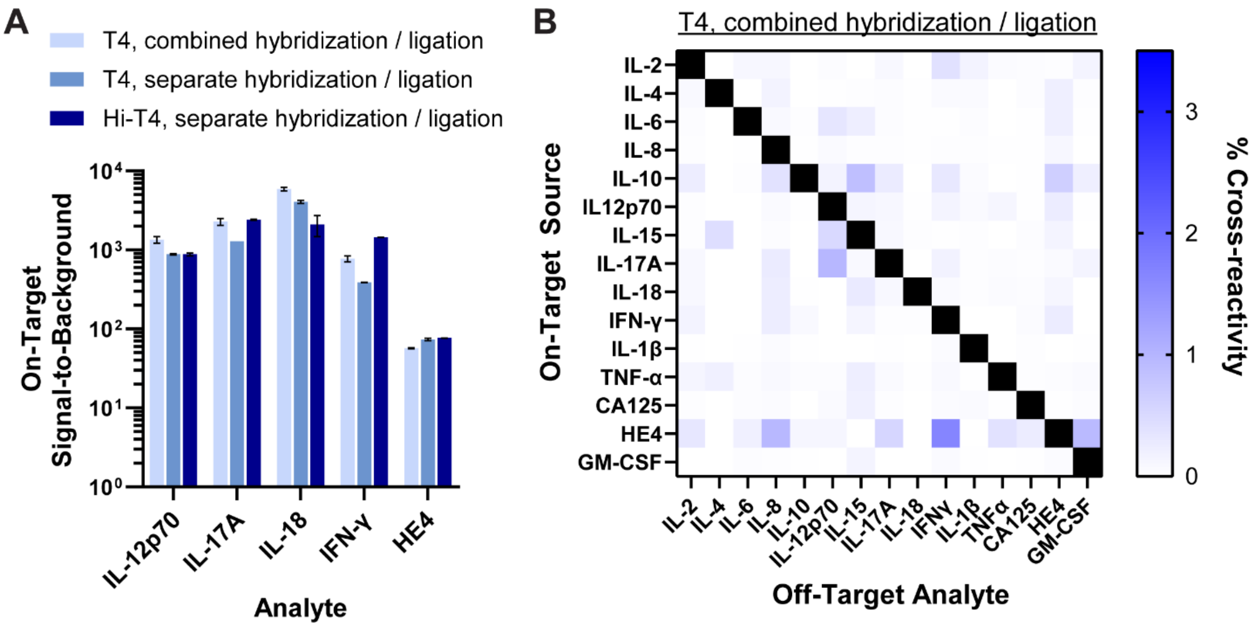


**Figure S8. Comparison of ligation conditions in 15-plex PRO-MOSAIX assay. (A)** Signal-to-background ratios for on-target beads in a four-protein mixture measured by the 15-plex PRO-MOSAIX assay with solution RCA, using (1) T4 DNA ligase with a combined connector oligo hybridization and ligation step performed at 37°C for 30 minutes; (2) T4 DNA ligase with separate hybridization and ligation steps, each performed at 37°C for 30 minutes; and (3) Hi-T4 DNA ligase with separate hybridization and ligation steps, performed at 37°C and 45°C, respectively, each for 30 minutes. Error bars represent the standard deviation of duplicate measurements. **(B)** Percent cross-reactivities in 15-plex PRO-MOSAIX assay with gel RCA (2400 µL gel), using T4 DNA ligase with a combined connector oligo hybridization and ligation step. Percent cross-reactivities were calculated as the ratio of off- to on-target bead (signal AMB - background AMB) values. The data represent the mean of duplicate measurements.

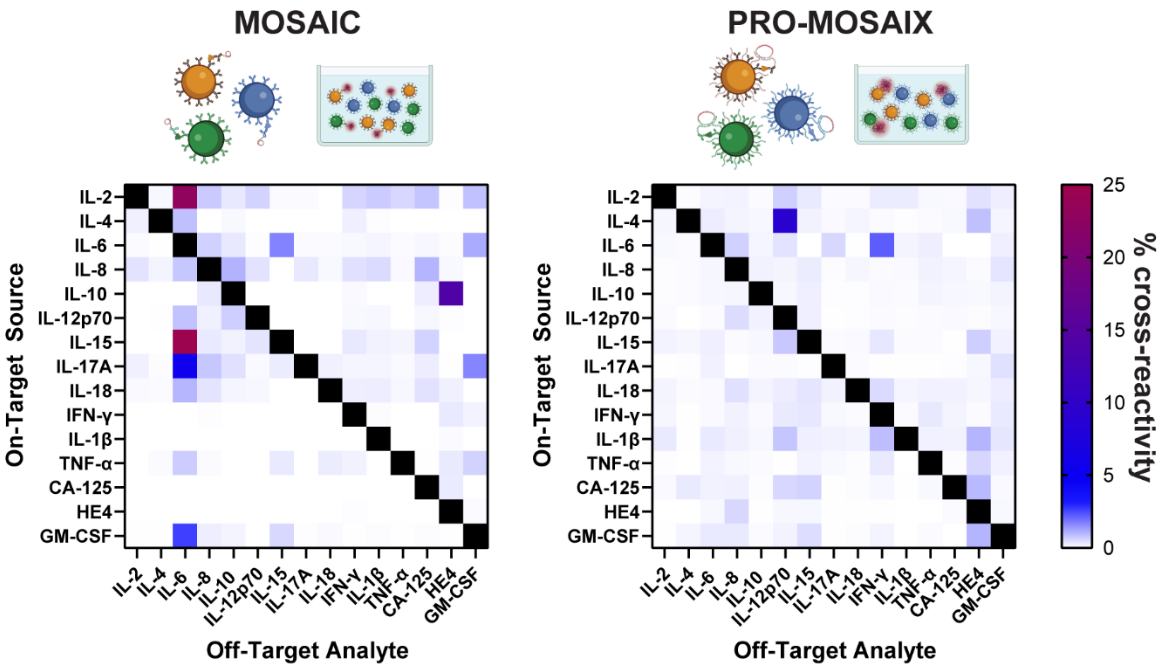


**Figure S9. Cross-reactivities in 15-plex MOSAIC and PRO-MOSAIX assays performed with solution RCA.** Single protein high-concentration samples (average on-target bead AMB values over 1) were measured. Percent cross-reactivities were calculated as the ratio of off- to on-target bead (signal AMB - background AMB) values. A gel volume of 2400 µL was used for RCA. The data represent the mean of duplicate measurements. The color gradient below 3.5% cross-reactivity was matched to be the same as the color gradient displayed for the MOSAIC and PRO-MOSAIX 15-plex assays with gel RCA in **Figure 5**.

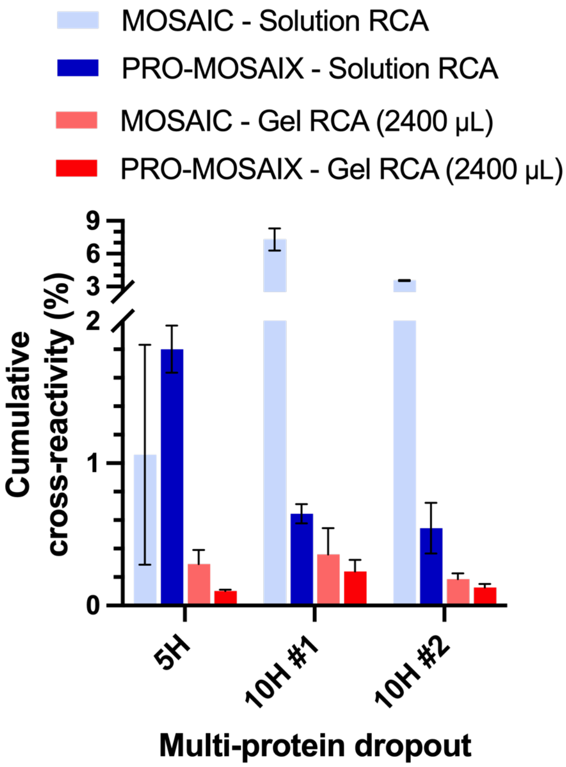


**Figure S10. Cumulative cross-reactivity in 15-plex MOSAIC and PRO-MOSAIX multi-protein dropout assays.** Measurements were performed with a 2400 μL gel volume for gel-based RCA. Light and cyan blue bars for solution-based MOSAIC and PRO-MOSAIX correspond to those shown in **Figure 5B**. Error bars represent the standard deviation of n=2-3 replicates.

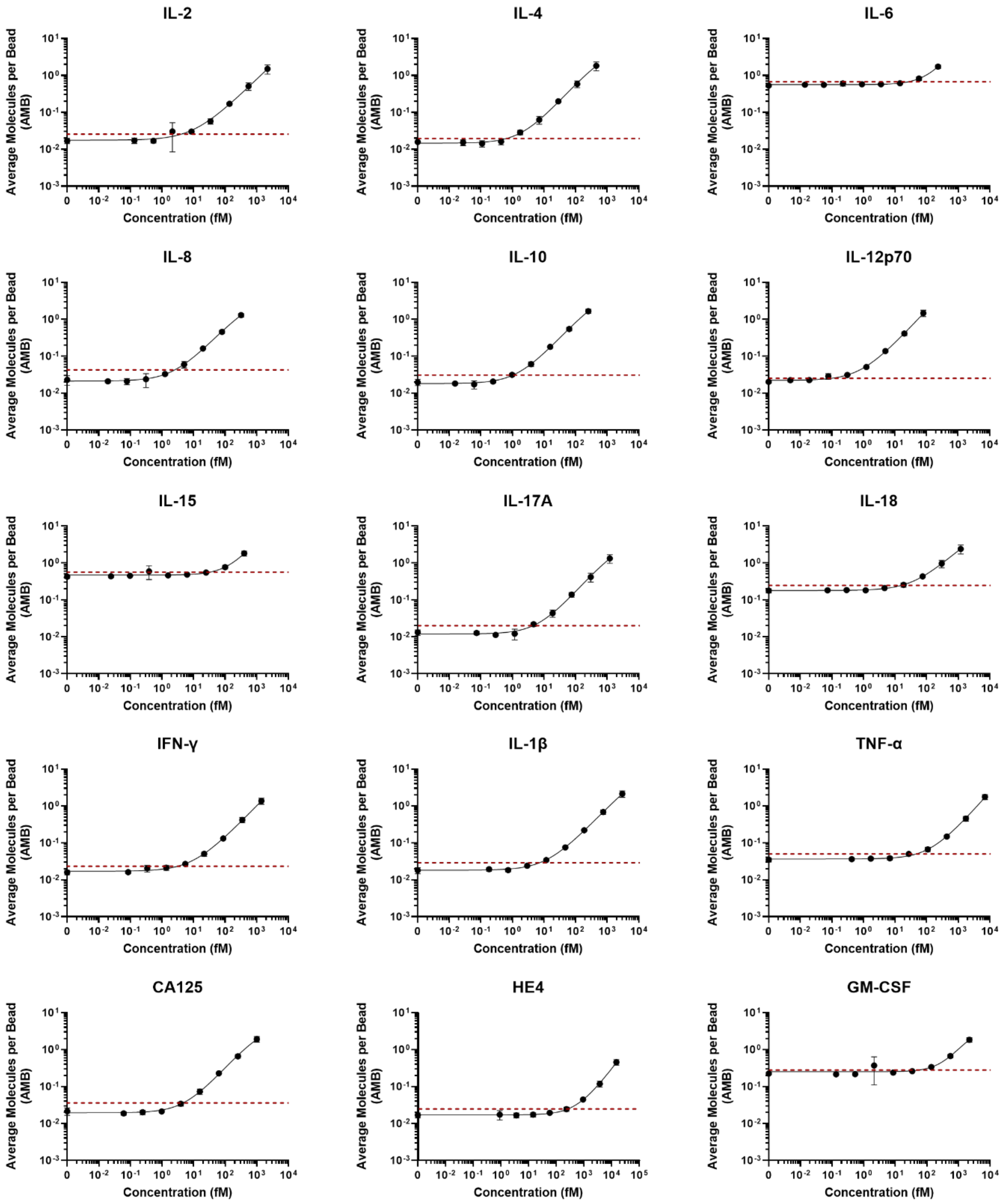


**Figure S11. Representative calibration curves for 15-plex MOSAIC assay performed with gel-based RCA.** Calibration curves for each analyte in the 15-plex MOSAIC assay are shown individually, with red dashed lines indicating the limit of detection (LOD). A gel volume of 2,400 μL was used for RCA. Error bars represent the standard deviation of triplicate measurements, with n=4 replicates for the blank. The LODs and LLOQs are summarized in **Table S3**.

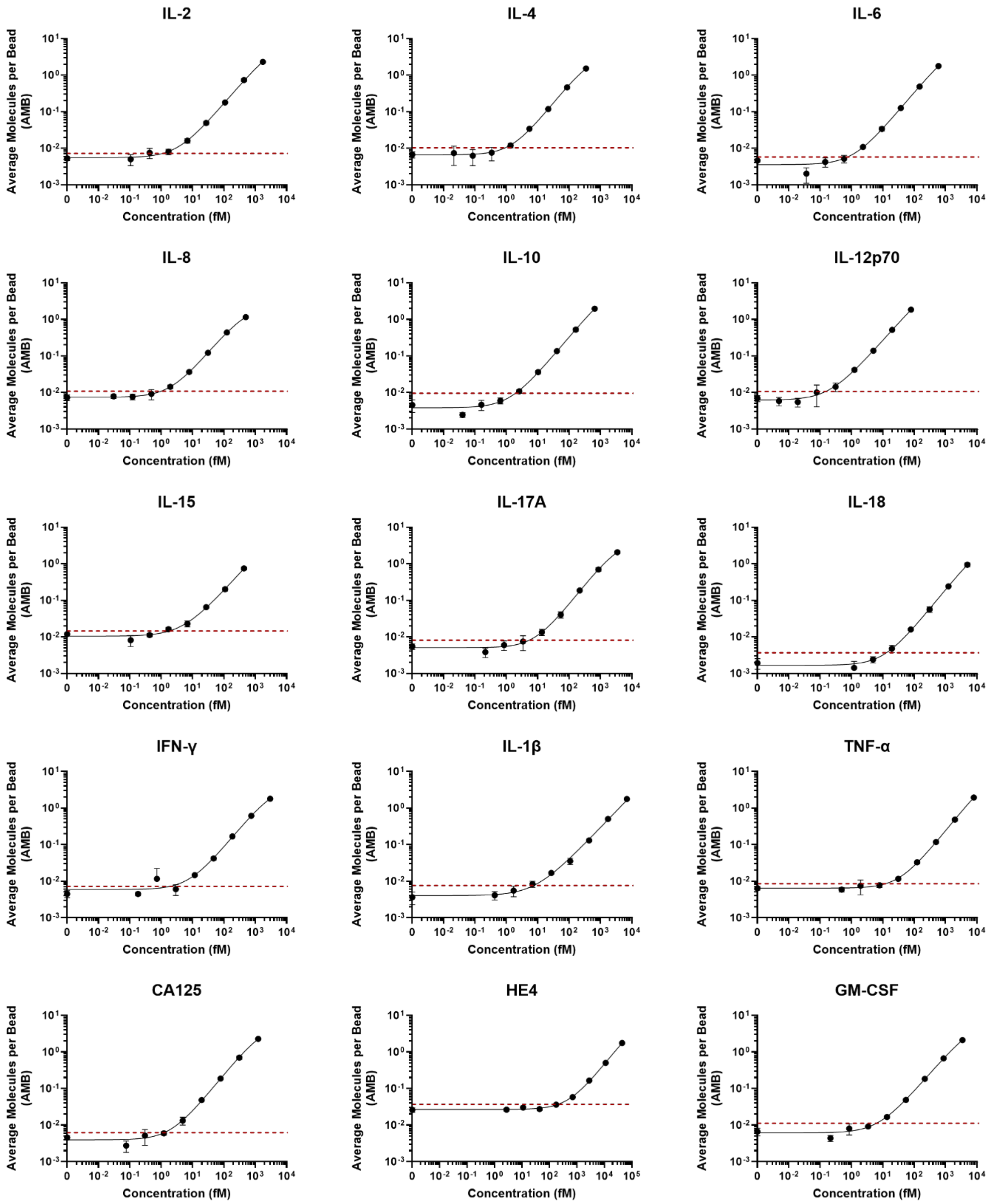


**Figure S12. Representative calibration curves for 15-plex PRO-MOSAIX assay performed with gel-based RCA.** Calibration curves for each analyte in the 15-plex PRO-MOSAIX assay are shown individually, with red dashed lines indicating the limit of detection (LOD). A gel volume of 2,400 μL was used for RCA. Error bars represent the standard deviation of triplicate measurements, with n=4 replicates for the blank. The LODs and LLOQs are summarized in **Table S3**.

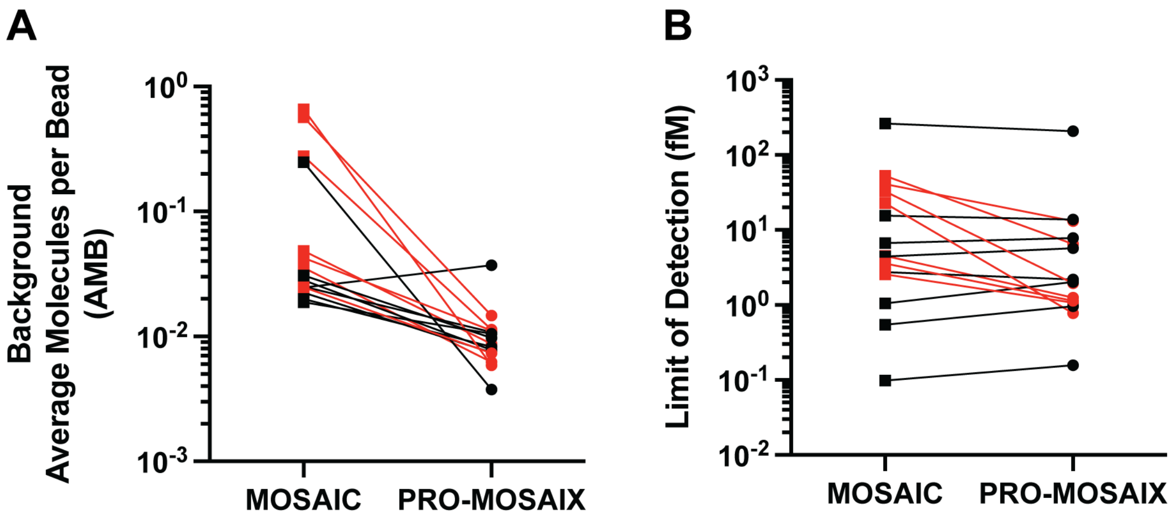


**Figure S13. Backgrounds and analytical sensitivities of 15-plex MOSAIC and PRO-MOSAIX assays with gel-based RCA. (A-B)** Background average molecules per bead (AMB) values **(A)** and limits of detection **(B)** for each analyte in 15-plex MOSAIC and PRO-MOSAIX assays. Elevated backgrounds are observed in MOSAIC assays compared to PRO-MOSAIX assays due to signal from cumulative nonspecific binding of detector antibodies. Red data points highlight analytes with elevated backgrounds and correspondingly lower sensitivities in MOSAIC compared to PRO-MOSAIX assays (IL-2, IL-6, IL-8, IL-15, TNF-α, CA125, and GM-CSF). A gel volume of 2,400 µL was used for RCA. The data represent the mean of n=3-4 replicates.

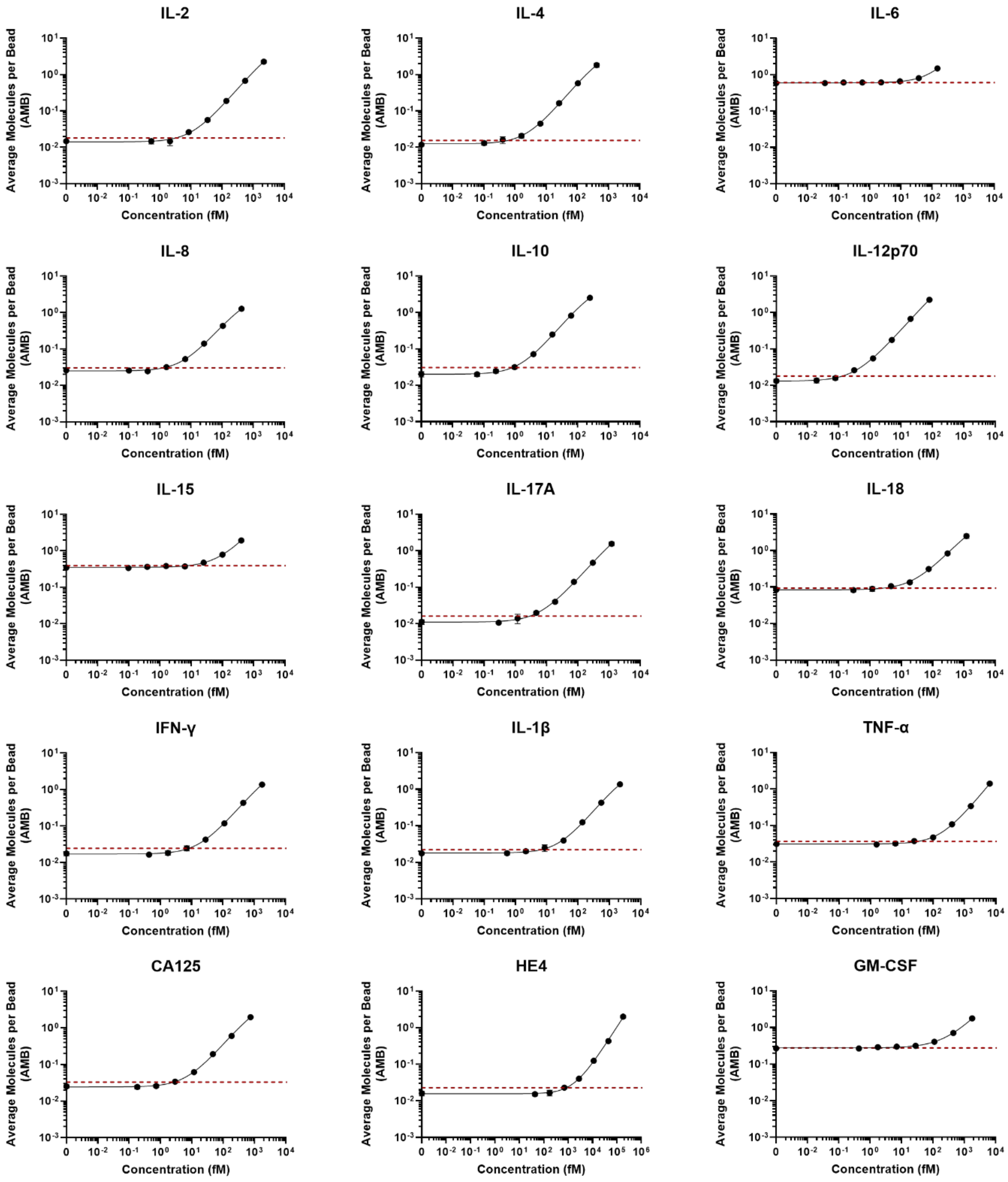


**Figure S14. Representative calibration curves for 15-plex MOSAIC assay used for human plasma measurements, performed with solution-based RCA.** Calibration curves for each analyte in the 15-plex MOSAIC assay are shown individually, with red dashed lines indicating the limit of detection (LOD). Error bars represent the standard deviation of triplicate measurements, with n=4 replicates for the blank. The LODs and LLOQs are summarized in **Table S4**.

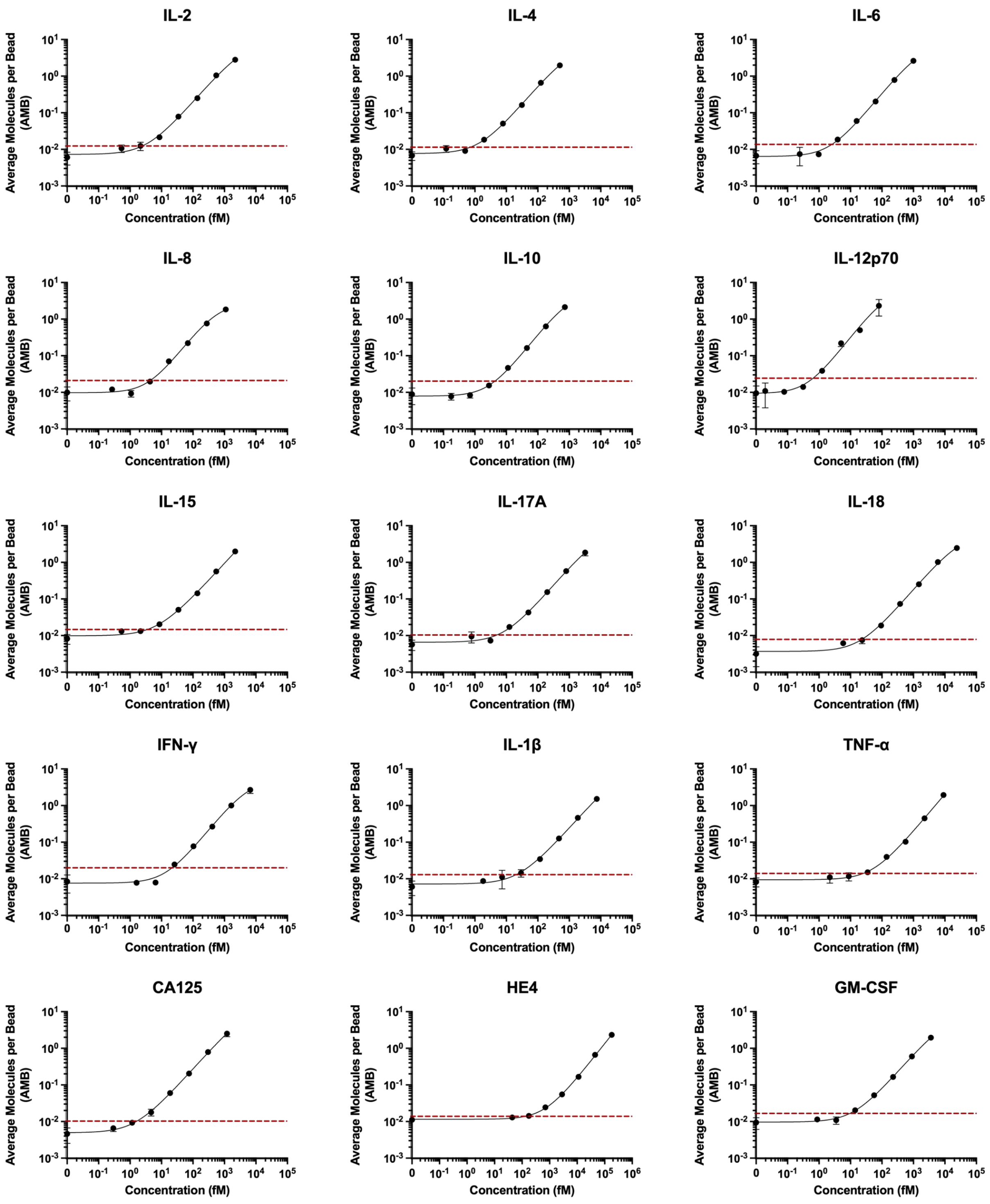


**Figure S15. Representative calibration curves for 15-plex PRO-MOSAIX assay used for human plasma measurements, performed with gel-based RCA.** Calibration curves for each analyte in the 15-plex PRO-MOSAIX assay are shown individually, with the red dashed line indicating the limit of detection (LOD). A gel volume of 1,600 μL was used for RCA. Error bars represent the standard deviation of triplicate measurements, with n=4 replicates for the blank. The LODs and LLOQs are summarized in **Table S5**.

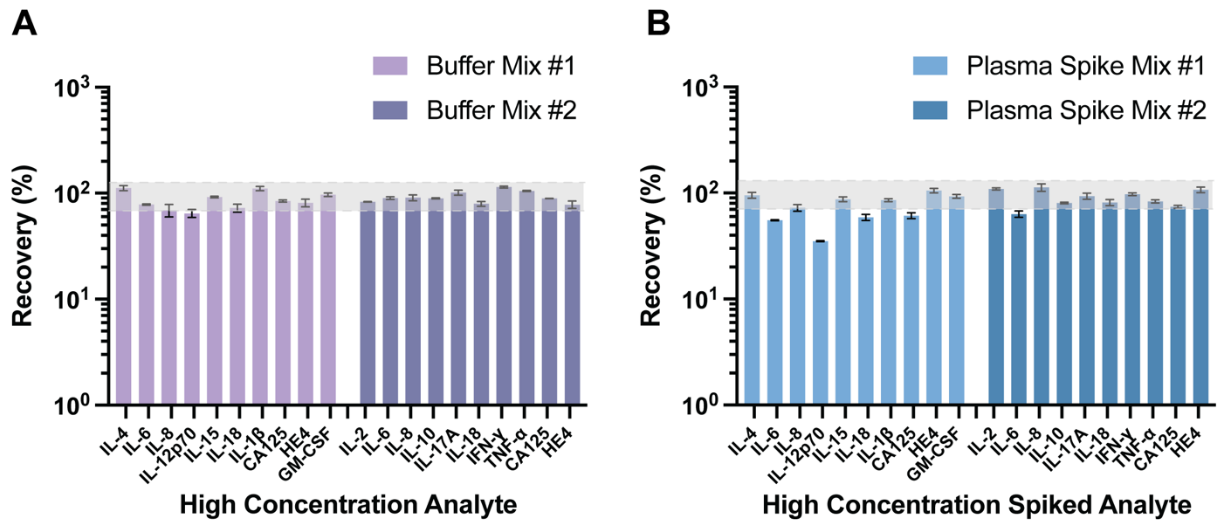


**Figure S16. Measurement accuracies of 15-plex PRO-MOSAIX assay for high concentration analytes in buffer and human plasma.** Recovery rates of 15-plex PRO-MOSAIX assay for high concentration analytes in recombinant protein mixtures in buffer **(A)** and spiked into 12-fold diluted pooled human plasma mixtures **(B)**. High concentrations were selected based on the upper end of the assay dynamic range. A gel volume of 1,600 µL gel was used for RCA. Recoveries were calculated as the ratio of measured to actual analyte or spike concentrations, with the acceptable 70–130% range shaded in gray. Analyte concentrations used are listed in **Table S6**. Error bars represent the standard deviation of n=2-3 replicates.

**Table S1. Amounts of fluorescent dye hydrazides used to make dye-encoded beads.** Amounts correspond to a starting number of $3.75\times10^8$ LodeStars carboxyl magnetic beads.

| Analyte | CF 488A Hydrazide (Biotium, #92152) | BDP 558/568 hydrazide (Lumiprobe, #17470) | Cyanine7 hydrazide (Lumiprobe, #15070) |
|---|---|---|---|
| **IL12-p70** | | | |
| **IL-10** | 5 µg | | |
| **IL-6** | 100 µg | | |
| **IFN-γ** | | 0.006 µg | |
| **IL-18** | | 0.1 µg | |
| **IL-8** | | | 0.025 µg |
| **IL-4** | | 0.1 µg | 0.025 µg |
| **CA125** | 100 µg | 0.006 µg | |
| **HE4** | 5 µg | 0.1 µg | |
| **TNF-α** | 100 µg | 0.1 µg | |
| **GM-CSF** | 5 µg | | 0.025 µg |
| **IL-1β** | 5 µg | 0.006 µg | |
| **IL-15** | 100 µg | | 0.025 µg |
| **IL-2** | 5 µg | 0.1 µg | 0.025 µg |
| **IL-17A** | 100 µg | 0.1 µg | 0.025 µg |

**Table S2. Analyte mixtures used in the multi-protein dropout experiment in Figure 5B**. High concentrations were selected based on the upper end of the assay dynamic range.

| Multi-protein dropout mixture | High Concentration Analyte |
|---|---|
| 5H | IL-4, IL-8, IL-18, CA125, HE4 |
| 10H #1 | IL-2, IL-4, IL-8, IL-10, IL-12p70, IL-15, IL-18, IFN-γ, CA125, HE4 |
| 10H #2 | IL-2, IL-6, IL-8, IL-10, IL-12p70, IL-17A, IFN-γ, TNF-α, CA125, GM-CSF |

**Table S3. Limit of detection (LOD) and lower limit of quantification (LLOQ) values of 15-plex MOSAIC and PRO-MOSAIX assay with gel RCA, using a gel volume of 2400 µL.** LOD and LLOQ values were determined as concentrations corresponding to three and ten standard deviations, respectively, above the background AMB.

| Analyte | MOSAIC | | PRO-MOSAIX | |
|---|---|---|---|---|
| | LOD (fM) | LLOQ (fM) | LOD (fM) | LLOQ (fM) |
| **IL-2** | 3.59 | 16.7 | 1.14 | 4.11 |
| **IL-4** | 0.546 | 1.53 | 0.959 | 2.86 |
| **IL-6** | 22.8 | 84.3 | 0.773 | 1.71 |
| **IL-8** | 2.56 | 9.13 | 1.08 | 3.54 |
| **IL-10** | 1.05 | 3.48 | 2.04 | 6.00 |
| **IL-12p70** | 0.098 | 0.542 | 0.157 | 0.480 |
| **IL-15** | 32.7 | 133 | 1.94 | 5.21 |
| **IL-17A** | 4.43 | 13.8 | 5.71 | 15.7 |
| **IL-18** | 15.6 | 66.4 | 13.6 | 39.3 |
| **IFN-γ** | 2.73 | 12.6 | 2.20 | 11.2 |
| **IL-1β** | 6.69 | 25.9 | 7.84 | 32.4 |
| **TNF-α** | 41.0 | 159.2 | 13.2 | 40.5 |
| **CA125** | 4.53 | 14.4 | 1.26 | 3.21 |
| **HE4** | 263 | 902 | 209 | 755 |
| **GM-CSF** | 53.0 | 247 | 6.48 | 20.0 |

**Table S4. Limit of detection (LOD) and lower limit of quantification (LLOQ) values of 15-plex MOSAIC and PRO-MOSAIX assay with solution RCA.** LOD and LLOQ values were determined as concentrations corresponding to three and ten standard deviations, respectively, above the background AMB. Values for PRO-MOSAIX represent the mean ± standard deviation [range] of two independent calibration curves performed on different days.

| Analyte | MOSAIC | | PRO-MOSAIX | |
|---|---|---|---|---|
| | LOD (fM) | LLOQ (fM) | LOD (fM) | LLOQ (fM) |
| **IL-2** | 3.16 | 9.10 | 0.722 ± 0.110 [0.644 – 0.800] | 2.53 ± 0.69 [2.05 – 3.02] |
| **IL-4** | 0.546 | 2.06 | 0.722 ± 0.024 [0.738 – 0.705] | 2.03 ± 0.04 [2.00 – 2.05] |
| **IL-6** | 1.41 | 9.11 | 0.704 ± 0.278 [0.508 – 0.901] | 2.29 ± 0.67 [1.81 – 2.76] |
| **IL-8** | 1.23 | 3.40 | 0.677 ± 0.111 [0.599 – 0.756] | 2.59 ± 0.39 [2.31 – 2.86] |
| **IL-10** | 0.800 | 2.53 | 0.260 ± 0.062 [0.304 – 0.216] | 0.682 ± 0.337 [0.443 – 0.920] |
| **IL-12p70** | 0.123 | 0.418 | 0.044 ± 0.028 [0.024 – 0.063] | 0.156 ± 0.075 [0.103 – 0.208] |
| **IL-15** | 7.77 | 31.4 | 2.71 ± 0.34 [2.47 – 2.96] | 8.93 ± 2.50 [7.16 – 10.70] |
| **IL-17A** | 2.87 | 9.55 | 2.35 ± 0.56 [1.95 – 2.74] | 7.79 ± 1.79 [6.52 – 9.05] |
| **IL-18** | 2.85 | 9.70 | 6.02 ± 3.86 [3.29 – 8.75] | 20.4 ± 12.7 [11.4 – 29.4] |
| **IFN-γ** | 8.15 | 25.4 | 13.7 ± 15.4 [2.8 – 24.5] | 37.3 ± 41.6 [7.8 – 66.7] |
| **IL-1β** | 6.46 | 21.6 | 18.3 ± 12.8 [9.3 – 27.3] | 53.4 ± 35.1 [28.6 – 78.3] |
| **TNF-α** | 35.3 | 111 | 15.2 ± 11.4 [7.1 – 23.2] | 41.7 ± 28.5 [21.5 – 61.9] |
| **CA125** | 2.67 | 8.05 | 0.510 ± 0.066 [0.464 – 0.557] | 1.65 ± 0.18 [1.52 – 1.77] |
| **HE4** | 875 | 2,695 | 319 ± 284 [118 – 520] | 1,166 ± 868 [553 – 1,780] |
| **GM-CSF** | 0.166 | 12.59 | 0.937 ± 0.971 [0.251 – 1.624] | 5.96 ± 3.03 [3.81 – 8.10] |

**Table S5. Limit of detection (LOD) and lower limit of quantification (LLOQ) values of 15-plex PRO-MOSAIX assay used in plasma measurements.** A gel volume of 1,600 µL was used for RCA. LOD and LLOQ values were determined as concentrations corresponding to three and ten standard deviations, respectively, above the background AMB.

| Analyte | PRO-MOSAIX | |
|---|---|---|
| | LOD (fM) | LLOQ (fM) |
| **IL-2** | 2.74 | 10.8 |
| **IL-4** | 0.827 | 3.15 |
| **IL-6** | 2.81 | 8.66 |
| **IL-8** | 4.85 | 14.3 |
| **IL-10** | 4.74 | 13.7 |
| **IL-12p70** | 0.708 | 2.07 |
| **IL-15** | 4.30 | 19.0 |
| **IL-17A** | 5.97 | 22.7 |
| **IL-18** | 25.9 | 93.4 |
| **IFN-γ** | 23.9 | 71.3 |
| **IL-1β** | 23.7 | 93.5 |
| **TNF-α** | 31.0 | 115 |
| **CA125** | 1.83 | 6.71 |
| **HE4** | 176 | 672 |
| **GM-CSF** | 10.8 | 37.1 |

**Table S6. Analyte concentrations used in the high/low-concentration protein mixtures and human plasma spike mixtures for the 15-plex PRO-MOSAIX assay in Figure 6A-B.**

| Analyte | Protein mixtures in buffer | | Human plasma spike mixtures | |
|---|---|---|---|---|
| | Buffer mix #1 (fM) | Buffer mix #2 (fM) | Plasma spike mix #1 (fM) | Plasma spike mix #2 (fM) |
| **IL-2** | 30 | 1,300 | 30 | 1,300 |
| **IL-4** | 300 | 10 | 380 | 10 |
| **IL-6** | 500 | 500 | 800 | 800 |
| **IL-8** | 660 | 660 | 600 | 600 |
| **IL-10** | 25 | 500 | 20 | 500 |
| **IL-12p70** | 55 | 2 | 60 | 4 |
| **IL-15** | 1200 | 35 | 1700 | 80 |
| **IL-17A** | 40 | 2,000 | 40 | 2,400 |
| **IL-18** | 14,000 | 14,000 | 20,000 | 20,000 |
| **IFN-γ** | 80 | 4,000 | 160 | 4,800 |
| **IL-1β** | 4,500 | 150 | 6,000 | 200 |
| **TNF-α** | 150 | 5,400 | 200 | 7200 |
| **CA125** | 720 | 720 | 900 | 900 |
| **HE4** | 12,000 | 12,000 | 45,000 | 45,000 |
| **GM-CSF** | 2,400 | 50 | 2,800 | 50 |

**Table S7. Recoveries of spiked mixtures of 15 medium- and high-concentration proteins in 12-fold diluted pooled human plasma, with the 15-plex PRO-MOSAIX assay.** A gel volume of 1,600 µL was used for RCA. Plasma was diluted using sample diluent consisting of 1x PBS with 2% BSA, 5 mM EDTA, 0.1% Tween-20, 0.1 mg/mL heparin (MilliporeSigma, H3393), and 0.15% ProClin 300. Recoveries are reported as the mean ± standard deviation of duplicate measurements.

| Analyte | Medium concentration mixture | | High concentration mixture | |
|---|---|---|---|---|
| | Spiked Protein (fM) | Recoveries (%) | Spiked Protein (fM) | Recoveries (%) |
| **IL-2** | 120 | 102.8 ± 8.2 | 1,200 | 108.2 ± 7.1 |
| **IL-4** | 30 | 92.7 ± 11.3 | 350 | 98.7 ± 2.3 |
| **IL-6** | 60 | 69.3 ± 7.9 | 720 | 59.2 ± 0.9 |
| **IL-8** | 60 | 113.4 ± 11.8 | 540 | 87.5 ± 0.5 |
| **IL-10** | 45 | 88.1 ± 3.7 | 450 | 80.3 ± 2.0 |
| **IL-12p70** | 20 | 66.2 ± 3.7 | 50 | 65.0 ± 0.0 |
| **IL-15** | 100 | 106.9 ± 10.6 | 1,500 | 96.4 ± 4.3 |
| **IL-17A** | 200 | 90.1 ± 2.7 | 2,100 | 91.9 ± 1.7 |
| **IL-18** | 1,000 | 97.6 ± 9.2 | 18,000 | 82.1 ± 4.3 |
| **IFN-γ** | 250 | 111.0 ± 0.4 | 4,300 | 90.5 ± 2.0 |
| **IL-1β** | 500 | 81.6 ± 1.5 | 5,400 | 73.1 ± 3.9 |
| **TNF-α** | 500 | 78.2 ± 8.4 | 6,400 | 80.6 ± 0.8 |
| **CA125** | 75 | 123.1 ± 22.3 | 800 | 72.7 ± 1.4 |
| **HE4** | 30,000 | 136.8 ± 15.2 | 40,000 | 90.4 ± 0.0 |
| **GM-CSF** | 220 | 84.2 ± 1.4 | 2,500 | 84.2 ± 3.1 |

**Table S8. Sequences of DNA oligos used in MOSAIC assays.** Bolded sequences correspond to the complementary regions in the primer and template. Red underlined sequence corresponds to the region in the circularized template encoding sequences for hybridization of ATTO 647N-labeled DNA probe. 5'-phos, 5' phosphate group; invdT, inverted dT.

| | |
|---|---|
| **Primer** | 5'-Azide- TTTTTTTTTTTTTTT**TAGACACCGTTCCTTGGACAGA*G*C** |
| **Template** | 5'-phos-**GAACGGTGTCTA**TTATGTCCTATCCTCAGC TATTATGTCCTATCCTCAGC TATTATGTCCTATCCTCA**GCTCTGTCCAAG** |
| **Probe** | 5'-[ATTO 647N]- TATTATGTCCTATCCTCAGC -invdT |

**Table S9. Sequences of DNA oligos used in PRO-MOSAIX assays.** Color-matched highlighted sequences correspond to hybridization regions of connector and proximity oligos. Red underlined sequences correspond to the region in the circularized template encoding sequences for hybridization of ATTO 647N-labeled DNA probe. 5'-phos: 5' phosphate group.

| | | |
|---|---|---|
| **IL-2** | Capture Proximity Oligo | 5'-NH2(C6)- AAAAAAAAAA CCTTGTTCACA TTGCTAGAC*C*T |
| | Detector Proximity Oligo | 5'-Azide- AAAAAAAAAA TCTACGGTATTTGA ACTATTCTG*C*T |
| | Connector Oligo 1 | 5'-phos- TCAAATACCGTAGA AAAAAA AGGTCTAGCAA |
| | Connector Oligo 2 | 5'-phos- TGTGAACAAGG TATTATGTCCTATCCTCAGC TATTATGTCCTATCCTCAGC AGCAGAATAGT |
| **IL-4** | Capture Proximity Oligo | 5'-NH2(C6)- AAAAAAAAAA GATCACATCCA ACGAAGTCA*T*C |
| | Detector Proximity Oligo | 5'-Azide- AAAAAAAAAA ACAGATAACGATTT GGAAGTCAA*C*T |
| | Connector Oligo 1 | 5'-phos- AAATCGTTATCTGT AAAAAA GATGACTTCGT |
| | Connector Oligo 2 | 5'-phos-TGGATGTGATCTATTATGTCCTATCCTCAGC TATTATGTCCTATCCTCAGC AGTTGACTTCC |
| **IL-6** | Capture Proximity Oligo | 5'-NH2(C6)- AAAAAAAAAA TCGATTACTCT AGTCTCTCT*C*A |
| | Detector Proximity Oligo | 5'-Azide- AAAAAAAAAA CAATATAGCTTTGA TGTAAGTCA*T*C |
| | Connector Oligo 1 | 5'-phos- TCAAAGCTATATTG AAAAAA TGAGAGAGACT |
| | Connector Oligo 2 | 5'-phos- AGAGTAATCGA TATTATGTCCTATCCTCAGC TATTATGTCCTATCCTCAGC GATGACTTACA |
| **IL-8** | Capture Proximity Oligo | 5'-NH2(C6)- AAAAAAAAAA GTTTCAAGTCC GACATGCTC*T*A |
| | Detector Proximity Oligo | 5'-Azide- AAAAAAAAAA ACATTTGTAGATCA ACAGTAATC*G*T |

| | | |
|---|---|---|
| | Connector Oligo 1 | 5'-phos- TGATCTACAAATGT AAAAAA TAGAGCATGTC |
| | Connector Oligo 2 | 5'-phos- GGACTTGAAAC TATTATGTCCTATCCTCAGC TATTATGTCCTATCCTCAGC ACGATTACTGT |
| **IL-10** | Capture Proximity Oligo | 5'-$NH_2$(C6)- AAAAAAAAAA CGAGAAGATCT TGTCAAATC*G*T |
| | Detector Proximity Oligo | 5'-Azide- AAAAAAAAAA TGACAATAAACAGG CATAGTTCT*C*A |
| | Connector Oligo 1 | 5'-phos- CTTGTTTATTGTCAAAAAAAACGATTTGACA |
| | Connector Oligo 2 | 5'-phos – AGATCTTCTCG TATTATGTCCTATCCTCAGC TATTATGTCCTATCCTCAGC TGAGAACTATG |
| **IL-12p70** | Capture Proximity Oligo | 5'-$NH_2$(C6)- AAATAAATAAATAAATAAAT GACGTTCTTACAGG GCAAGAGTATTA*C*C |
| | Detector Proximity Oligo | 5'-Azide- AAAAAAAAAA GAATATGACAGAGC GGTTAGACACTA*C*G |
| | Connector Oligo 1 | 5'-phos- GCTCTGTCATATTCAAAAAAGGTAATACTCTTGC |
| | Connector Oligo 2 | 5'-phos- CCTGTAAGAACGTC TATTATGTCCTATCCTCAGC TATTATGTCCTATCCTCAGC CGTAGTGTCTAACC |
| **IL-15** | Capture Proximity Oligo | 5'-$NH_2$(C6)- AAAAAAAAAA CAGAAATTGCA TCAAATGCT*C*T |
| | Detector Proximity Oligo | 5'-Azide- AAAAAAAAAA AGTTCAACTGTAGA ATCTTCATC*G*A |
| | Connector Oligo 1 | 5'-phos- TCTACAGTTGAACT AAAAAA AGAGCATTTGA |
| | Connector Oligo 2 | 5'-phos- TGCAATTTCTG TATTATGTCCTATCCTCAGC TATTATGTCCTATCCTCAGC TCGATGAAGAT |
| **IL-17A** | Capture Proximity Oligo | 5'-$NH_2$(C6)- AAAAAAAAAA AACGTCATACC ACAACCTAA*C*C |
| | Detector Proximity Oligo | 5'-Azide- AAAAAAAAAA GATGACCTAGTTTG CGTTAAACA*C*T |
| | Connector Oligo 1 | 5'-phos- CAAACTAGGTCATC AAAAAA GGTTAGGTTGT |
| | Connector Oligo 2 | 5'-phos- GGTATGACGTT TATTATGTCCTATCCTCAGC TATTATGTCCTATCCTCAGC AGTGTTTAACG |
| **IL-18** | Capture Proximity Oligo | 5'-$NH_2$(C6)- AAAAAAAAAA CCATGTCAATG ACTTTGAAT*G*C |
| | Detector Proximity Oligo | 5'-Azide- AAAAAAAAAA TGAACGAAAAATGC CAATCCTGT*C*T |

| | | |
|---|---|---|
| | Connector Oligo 1 | 5'-phos- GCATTTTTCGTTCA AAAAAA GCATTCAAAGT |
| | Connector Oligo 2 | 5'-phos- CATTGACATGG TATTATGTCCTATCCTCAGC TATTATGTCCTATCCTCAGC AGACAGGATTG |
| **IFN-γ** | Capture Proximity Oligo | 5'-$NH_2$(C6)- AAAAAAAAAA AGATTTCCAGG GTTCAAGTT*C*A |
| | Detector Proximity Oligo | 5'-Azide- AAAAAAAAAA ACGACAAGATTAGA GCCACTTAT*T*A |
| | Connector Oligo 1 | 5'-phos- TCTAATCTTGTCGT AAAAAA TGAACTTGAAC |
| | Connector Oligo 2 | 5'-phos- CCTGGAAATCT TATTATGTCCTATCCTCAGC TATTATGTCCTATCCTCAGC TAATAAGTGGC |
| **IL-1β** | Capture Proximity Oligo | 5'-$NH_2$(C6)- AAAAAAAAAA TCAACTTCTGT CGTTTCACA*T*G |
| | Detector Proximity Oligo | 5'-Azide- AAAAAAAAAA GAATCTGAACTTGT ATACGTTTG*A*C |
| | Connector Oligo 1 | 5'-phos- ACAAGTTCAGATTC AAAAAA CATGTGAAACG |
| | Connector Oligo 2 | 5'-phos- ACAGAAGTTGA TATTATGTCCTATCCTCAGC TATTATGTCCTATCCTCAGC GTCAAACGTAT |
| **TNF-α** | Capture Proximity Oligo | 5'-$NH_2$(C6)- AAAAAAAAAA ATGGAAACTGG AACAGTATG*G*C |
| | Detector Proximity Oligo | 5'-Azide- AAAAAAAAAA AATCACAATCACAG ACCTCTGAT*G*A |
| | Connector Oligo 1 | 5'-phos- CTGTGATTGTGATT AAAAAA GCCATACTGTT |
| | Connector Oligo 2 | 5'-phos- CCAGTTTCCAT TATTATGTCCTATCCTCAGC TATTATGTCCTATCCTCAGC TCATCAGAGGT |
| **CA125** | Capture Proximity Oligo | 5'-$NH_2$(C6)- AAAAAAAAAA CAACGAAAGGA ATTCTGCCA*A*T |
| | Detector Proximity Oligo | 5'-Azide- AAAAAAAAAA TAGGTAAACCAAAG GTCCATACG*A*A |
| | Connector Oligo 1 | 5'-phos- CTTTGGTTTACCTA AAAAAA ATTGGCAGAAT |
| | Connector Oligo 2 | 5'-phos- TCCTTTCGTTG TATTATGTCCTATCCTCAGC TATTATGTCCTATCCTCAGC TTCGTATGGAC |
| **HE4** | Capture Proximity Oligo | 5'-$NH_2$(C6)- AAAAAAAAAA TAATGGCAGAC TGTACGAGA*A*C |
| | Detector Proximity Oligo | 5'-Azide- AAAAAAAAAA TACAACAACATCAG TAGGTCCAA*C*T |

| | | |
|---|---|---|
| | Connector Oligo 1 | 5'-phos- CTGATGTTGTTGTA AAAAAA GTTCTCGTACA |
| | Connector Oligo 2 | 5'-phos- GTCTGCCATTA TATTATGTCCTATCCTCAGC TATTATGTCCTATCCTCAGC AGTTGGACCTA |
| GM-CSF | Capture Proximity Oligo | 5'-$NH_2$(C6)- AAAAAAAAAA CACGAGAGATT ACGTTGGTA*A*C |
| | Detector Proximity Oligo | 5'-Azide- AAAAAAAAAA CAAGCAATATGTGA CTTACAGTC*C*T |
| | Connector Oligo 1 | 5'-phos- TCACATATTGCTTG AAAAAA GTTACCAACGT |
| | Connector Oligo 2 | 5'-phos- AATCTCTCGTG TATTATGTCCTATCCTCAGC TATTATGTCCTATCCTCAGC AGGACTGTAAG |

**Table S10. Antibodies and recombinant protein standards used in this work.**

| Analyte | Capture Antibody | | Detector Antibody | | Recombinant Protein | |
|---|---|---|---|---|---|---|
| | Vendor | Catalog # | Vendor | Catalog # | Vendor | Catalog # |
| **IL-2** | R&D Systems | MAB602 | R&D Systems | MAB202 | R&D Systems | BT-002-010 |
| **IL-4** | BioLegend | 500702 | BioLegend | 500802 | R&D Systems | 204-IL-010 |
| **IL-6** | R&D Systems | MAB206 | R&D Systems | AF-206-NA | R&D Systems | 206-IL-010 |
| **IL-8** | BD Biosciences | 554716 | BD Biosciences | 550419 | R&D Systems | 208IL010 |
| **IL-10** | BioLegend | 506802 | BioLegend | 501504 | R&D Systems | 217-IL-005 |
| **IL-12p70** | BioLegend | 511002 | BioLegend | 508807 | R&D Systems | 219-IL-005 |
| **IL-15** | R&D Systems | MAB647 | R&D Systems | MAB247 | R&D Systems | 247-ILB-005 |
| **IL-17A** | R&D Systems | MAB317 | R&D Systems | AF-317-NA | BioLegend | 570509 |
| **IL-18** | R&D Systems | D044-3 | R&D Systems | D045-3 | R&D Systems | 9124-IL-010 |
| **IFN-γ** | BioLegend | 507502 | R&D Systems | MAB285 | R&D Systems | 285-IF-025 |
| **IL-1β** | BioLegend | 508202 | BioLegend | anti-human IL-1β clone H1b-98 | R&D Systems | 201-LB-005 |
| **TNF-α** | R&D Systems | MAB610 | Abcam | ab9635 | R&D Systems | 210-TA-005 |
| **CA125** | R&D Systems | DY5609-05 | R&D Systems | AF62981 | R&D Systems | DY5609-05 |
| **HE4** | R&D Systems | DY6274-05 | R&D Systems | AF6274 | R&D Systems | DY6274-05 |
| **GM-CSF** | R&D Systems | MAB615 | R&D Systems | MAB215 | R&D Systems | 215-GM-010 |

**Table S11. Conditions for antibody coupling to MOSAIC capture beads.** The EDC activation and antibody coupling steps were carried out at 4 °C for all beads.

| Analyte | Starting Bead Number | EDC (µL of 10 mg/mL solution) | Capture Antibody (µg) |
|---|---|---|---|
| **IL-2** | $3.0 \times 10^8$ | 8 | 43 |
| **IL-4** | $3.0 \times 10^8$ | 8 | 38 |
| **IL-6** | $2.0 \times 10^8$ | 4 | 25 |
| **IL-8** | $3.0 \times 10^8$ | 3 | 25 |
| **IL-10** | $2.8 \times 10^8$ | 3 | 25 |
| **IL-12p70** | $4.2 \times 10^8$ | 9 | 60 |
| **IL-15** | $3.0 \times 10^8$ | 8 | 43 |
| **IL-17A** | $3.0 \times 10^8$ | 8 | 43 |
| **IL-18** | $3.0 \times 10^8$ | 6 | 43 |
| **IFN-γ** | $3.0 \times 10^8$ | 6 | 40 |
| **IL-1β** | $3.0 \times 10^8$ | 6 | 43 |
| **TNF-α** | $3.0 \times 10^8$ | 6 | 43 |
| **CA125** | $3.0 \times 10^8$ | 6 | 43 |
| **HE4** | $3.0 \times 10^8$ | 6 | 43 |
| **GM-CSF** | $3.0 \times 10^8$ | 8 | 43 |

**Table S12. Conditions for antibody and DNA coupling to PRO-MOSAIX capture beads.** The EDC activation and antibody coupling steps were carried out at 4 °C for all beads. The mTz-DNA coupling to TCO/antibody-conjugated beads was carried out at room temperature.

| Analyte | Starting Bead Number | EDC (µL of 10 mg/mL solution) | Capture Antibody (µg) | TCO-PEG6-$NH_2$ (µg) | mTz-DNA (µM) |
|---|---|---|---|---|---|
| **IL-2** | $3.0 \times 10^8$ | 8 | 43 | 25 | 10 |
| **IL-4** | $3.0 \times 10^8$ | 8 | 41 | 25 | 10 |
| **IL-6** | $3.0 \times 10^8$ | 8 | 43 | 25 | 10 |
| **IL-8** | $3.0 \times 10^8$ | 5 | 20 | 25 | 10 |
| **IL-10** | $3.0 \times 10^8$ | 8 | 30 | 25 | 10 |
| **IL-12p70** | $3.0 \times 10^8$ | 8 | 43 | 25 | 10 |
| **IL-15** | $3.0 \times 10^8$ | 8 | 43 | 25 | 10 |
| **IL-17A** | $3.0 \times 10^8$ | 8 | 43 | 25 | 10 |
| **IL-18** | $3.0 \times 10^8$ | 8 | 43 | 25 | 10 |
| **IFN-γ** | $3.0 \times 10^8$ | 8 | 43 | 25 | 10 |
| **IL-1β** | $3.0 \times 10^8$ | 8 | 43 | 25 | 10 |
| **TNF-α** | $3.0 \times 10^8$ | 8 | 43 | 25 | 10 |
| **CA125** | $3.0 \times 10^8$ | 8 | 43 | 25 | 10 |
| **HE4** | $3.0 \times 10^8$ | 8 | 43 | 25 | 10 |
| **GM-CSF** | $3.0 \times 10^8$ | 8 | 43 | 25 | 10 |

**Table S13. Assay conditions for multiplex MOSAIC assays.** Solution-based RCA reaction times were 20 minutes. Gel-based RCA reaction times were 90 minutes, followed by melting, washing, and a 20-minute incubation with ATTO647N-DNA probe at 37°C for fluorescent labeling.

| Assay | Assay Type [a] | Incubation Times (minutes)<br>Sample / Detector | # Beads per Analyte | Detector Antibody (µg/mL) |
|---|---|---|---|---|
| **3-plex** | 2-step | 60 | 30,000 | 0.1 / 0.25 / 0.1<br>IL-12p70 / IL-10 / IL-6 |
| **15-plex** | 3-step | 60-15 | 20,000 | 0.1 each [b] |

[a] 2-step assays consist of sample incubation with both capture beads and detector, followed by washing and rolling circle amplification (RCA). 3-step assays consist of sample incubation with capture beads, washing, and incubation with detector, before washing and subsequent RCA.
[b] 0.05 µg/mL IL-8 and 0.025 µg/mL HE4 detector were used for the calibration curve with gel-based RCA.

**Table S14. Assay conditions for PRO-MOSAIX assays.** Solution-based RCA reaction times were 20 minutes. Gel-based RCA reaction times were 90 minutes, followed by melting, washing, and a 20-minute incubation with ATTO647N-DNA probe at 37°C for fluorescent labeling.

| Assay | Assay Type [a] | Incubation Times (minutes)<br>Sample / Detector - Hybridization / Ligation [b] | # Beads per Analyte | Detector Antibody (µg/mL) |
|---|---|---|---|---|
| **3-plex** | 2-step | 60-30 | 30,000 | 0.1 / 0.25 / 0.1<br>IL-12p70 / IL-10 / IL-6 |
| **15-plex** | 3-step | 60-15-30-30 | 20,000 | 0.1 each [c] |

[a] 2-step assays consist of sample incubation with both capture beads and detector, followed by washing and subsequent hybridization and ligation steps before rolling circle amplification (RCA). 3-step assays consist of sample incubation with capture beads, washing, and incubation with detector, before washing and subsequent hybridization, ligation, and RCA steps.
[b] 3-plex assays: combined hybridization and ligation steps at 37°C, using T4 DNA ligase. 15-plex assays: separate hybridization and ligation steps at 37°C and 45°C, respectively, using Hi-T4 DNA ligase.
[c] 0.025 µg/mL HE4 detector was used for protein dropout measurements. For calibration curves performed with 2,400 µL gel volume, 0.05 µg/mL IL-8 and 0.025 µg/mL HE4 detector concentrations were used. For measurements of plasma and the corresponding calibration curves performed with 1,600 µL gel volumes, 0.05 µg/mL IL-8 and and 0.007 µg/mL HE4 detector concentrations were used.

**Table S15. Rolling circle amplification (RCA) conditions for multiplex MOSAIC and PRO-MOSAIX assays.**

| | Solution RCA | | Gel RCA |
|---|---|---|---|
| | **Single-plex assays [a]** | **Optimized for multiplex assays** | **Multiplex MOSAIC and PRO-MOSAIX assays** |
| ***RCA reaction time*** | *45 minutes* | *20 minutes* | *90 minutes* |
| **phi29 DNA polymerase** (Biosearch Technologies) | 0.1 U/µL | 0.1 U/µL | 0.1 U/µL |
| **ATTO647N DNA probe** | 0.5 nM | 2 nM | -- |
| **10X Reaction Buffer** 500 mM Tris-HCl pH 7.5, 100 mM $MgCl_2$, 100 mM $(NH_4)_2SO_4$ | 1X | 1X | 1X |
| **dNTP Solution Mix** | 0.25 mM | 0.25 mM | 0.25 mM |
| **Bovine serum albumin** | 2 mg/mL | 2 mg/mL | 0.2 mg/mL |
| **Tween-20** | 0.2% | 0.2% | 0.2% |
| **Trehalose** | 0.1% | 0.1% | 0.1% |
| **NaCl** | - | 50 mM | - |
| **Heparin** | -- | 0.005 µg/mL | -- |

[a] Unoptimized solution RCA conditions for spike-in RCA control experiment in Figure S1 used 60-minute RCA reaction time and 0.5 mM dNTP solution mix.
